# Multilayer analysis reveals extensive hidden population complexity in a dairy starter microbiome comprising few species

**DOI:** 10.64898/2026.10.09.757929

**Authors:** Axel Soto-Serrano, Eoghan Reilly, Kelsey White, Wenwen Li, Shirin Safaeian, Jennifer Mahony, Paulina Deptula, Lukasz Krych

## Abstract

The genomic and epigenomic complexity of microbiomes with low species richness remains poorly characterized. Here, we integrated 111 non-redundant isolate genomes with deep long-read metagenomics to investigate an undefined mesophilic dairy starter. Closely related isolates harboured distinct combinations of features associated with fermentation traits and phage defence, including surface polysaccharide genotypes, plasmids, prophages, defence systems, and methylation profiles. Metagenomics revealed additional elements and inferred host associations. However, the limited overlap between approaches across core-genome lineages and accessory repertoires underscored complementary but incomplete recovery, to the extent that the dominant metagenomic cell wall polysaccharide genotype (29.6% abundance) was absent from isolates. Metagenomic methylation signatures supported 36 species-level plasmid–host assignments, corroborated by isolate genomes in nine of ten testable cases. Our findings revealed extensive combinatorial diversity hidden from either approach alone, providing a plausible molecular basis for phage resilience, while likely representing only a lower bound on population complexity.

## Introduction

Microbial communities are increasingly being studied at enhanced genome resolution, driven by rapid advances in sequencing technologies, read accuracy, and metagenome assembly methods. Long-read sequencing has substantially improved the recovery of near-complete bacterial chromosomes, strain-resolved variation, and circular mobile genetic elements (MGEs) directly from mixed DNA, while also enabling epigenetic profiling through native DNA methylation signals [1, 2]. In parallel, microbial cultivation followed by whole-genome sequencing (WGS) remains essential to assemble validated complete genomes of the highest quality, improving functional annotation [3], and enabling genotype–phenotype studies and experimental validation.

Despite recent advances, neither approach provides an unbiased representation of the genomic diversity present within a microbial community. Isolation-based approaches are subject to cultivation and sampling biases, which can underrepresent rare or slow growing lineages and overrepresent clonal selected strains favoured by the growth conditions [4]. Conversely, metagenome assembly and binning can systematically compromise the recovery of genetic features that are repetitive, present in closely related strains, variably abundant, or mobile across hosts [5, 6]. MGEs such as plasmids represent a good instance of these challenges, as they can share conserved backbones, undergo recombination, and occur at variable copy numbers that impair their abundance from host chromosomes, hindering both assembly and host attribution [6, 7].

These limitations are especially acute in microbiomes with low species richness. Closely related strains often have similar growth requirements, limiting their selective recovery by cultivation. At the same time, ecological and functional diversity in these communities may depend disproportionately on variation among strains with high genomic similarity. Much of this variation can be concentrated in MGEs [8], which mediate gene flux and often encode traits related to metabolism, stress tolerance, and phage defence [9–11]. In parallel, phage–host interaction determinants such as cell surface structures, prophages, and antiphage systems can also vary extensively among closely related strains [12, 13], reflecting the pervasive role of bacteriophages in shaping microbial population structure and community dynamics. Consequently, low-species, high-strain microbiomes are particularly challenging to resolve and provide an informative system for evaluating the complementary resolution of culture-dependent and culture-independent approaches.

Undefined mesophilic dairy starter cultures used for traditional Northern European cheeses, including Gouda, Cheddar and Danbo, provide a compelling example of this type of community. These consortia are typically described as complex communities dominated by *Lactococcus cremoris* and *Lactococcus lactis*, often accompanied by *Leuconostoc* species, and which retain elevated strain diversity that underpins fermentation performance, flavour development and resilience to bacteriophage predation [14, 15]. However, the extent of this complexity and the genomic layers through which it is structured remain unresolved. Importantly, many members of starter cultures are, in principle, readily culturable, enabling deep isolate sequencing and direct validation of community-level inferences. This combination of low species richness, extensive strain heterogeneity, and high cultivability, makes starter cultures a powerful model system to investigate how different genomic layers shape community complexity and how completely this diversity is captured by culture-dependent and culture-independent approaches.

Here, we use Tistrup, an undefined Danish mesophilic dairy starter culture maintained for more than a century, as a genome-resolved model of a low-species, high-strain microbiome. We combine deep long-read metagenomics with cultivation and WGS of 111 non-redundant strains to quantify overlap and complementarity between the two approaches across multiple, biologically relevant genomic layers. First, we resolve community composition and core-genome population structure and quantify the complementarity of metagenomics and isolate sequencing in recovering lineages and gene content. We then dissect below-species variation in cell wall polysaccharides (CWPS), exopolysaccharides (EPS), prophages, antiphage systems, and methylation repertoires, before characterizing plasmid diversity and leveraging native methylation signatures to infer plasmid–host associations directly from the metagenome [7, 16] (Fig. 1a). The integration of these genomic layers and DNA methylation reveals extensive hidden community complexity at different levels of resolution and shows that cultivation and metagenomics recover distinct fractions of the population, with substantial variation remaining inaccessible even when both approaches are combined.

**Figure 1.**
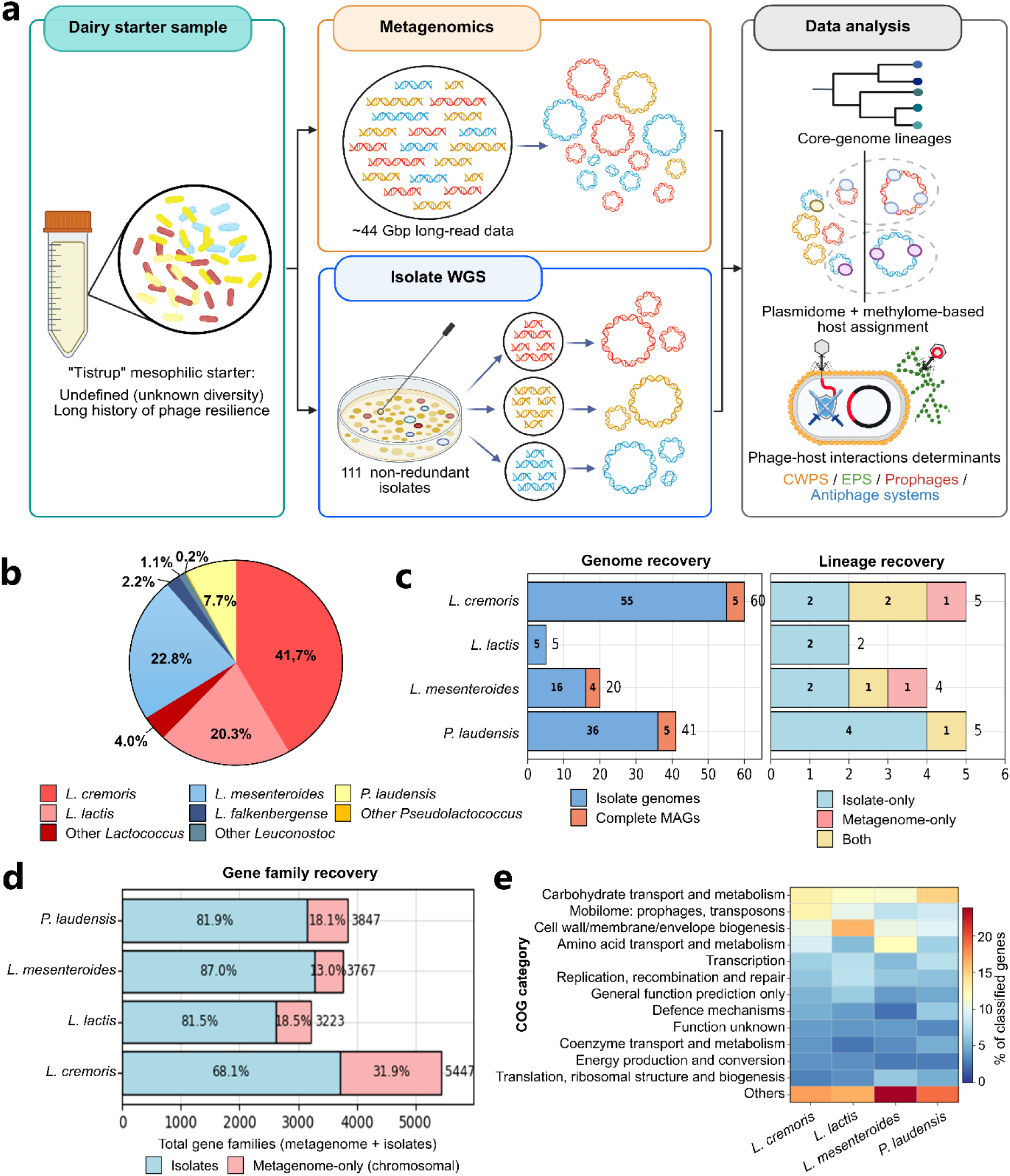
Community composition and complementary recovery of genomic diversity in Tistrup. **a** Experimental workflow combining long-read metagenomics with isolate whole-genome sequencing (WGS) to characterize genomic and epigenomic diversity. **b** Read-level taxonomic composition inferred using Metabuli, with additional taxa grouped by genus. **c** Recovery of isolate genomes and high-quality, 0% strain heterogeneity MAGs (left) and core-genome lineages (right) across the four cultivated species. Lineages are classified as isolate-only, metagenome-only or shared between approaches. Numbers within bars indicate category counts, and bar-end labels show totals. **d** Gene-family recovery from isolate genomes and a curated chromosomal metagenome catalogue excluding plasmid contigs. Internal labels indicate percentages of the combined gene-family pool within each species, while bar-end labels show total family counts. **e** Functional composition of metagenome-exclusive gene families assigned to Clusters of Orthologous Groups (COG) categories. Colours indicate the percentage of COG-assigned families belonging to each category within each species; families without a COG assignment are excluded from the calculations. “Others” combines categories not displayed individually. CWPS, cell wall polysaccharides; EPS, exopolysaccharides.

## Results

### Deep metagenomics and isolate sequencing reveal complementary population diversity

To characterize the composition and below-species diversity of the Tistrup starter culture, we combined deep long-read sequencing of both the whole mixed community and an extensive isolate collection (Figure 1a). For metagenome sequencing, we retained 3,934,275 reads comprising ∼44 Gb of sequence data (read N50 = 15,914 bp; median Q score = 24.4) for downstream analyses. Read-level taxonomic profiling revealed a community dominated by *L. cremoris*, followed by *Leuconostoc mesenteroides*, *L. lactis*, *Pseudolactococcus laudensis*, and *Leuconostoc falkenbergense*, while the remaining reads were assigned to closely related taxa within these genera (Fig. 1b). Metagenome assembly and binning yielded 27 metagenome-assembled genomes (MAGs), all taxonomically assigned to these same species. Fifteen MAGs were classified as high quality (>90% completeness; <5% contamination), whereas 12 MAGs were of medium quality (>50% completeness; <10% contamination) [17] (Fig. 1c; Table 1).

**Table 1.** Summary of MAGs recovered in this study.

| MAG ID | taxonomy | completeness,<br>contamination<br>(checkm2) | strain<br>heterogeneity<br>(checkm1) | contigs | longest<br>contig | genome<br>size | MIMAG<br>classification |
| --- | --- | --- | --- | --- | --- | --- | --- |
| bin.s.19 | *L. cremoris* | 100, 0.25 | 0 | 1 | 2412279 | 2412279 | High-quality |
| bin.s.20 | *L. cremoris* | 100, 0.25 | 0 | 1 | 2412184 | 2412184 | High-quality |
| bin.2.11 | *L. cremoris* | 66.37, 0.12 | 0 | 1 | 1626883 | 1626883 | Medium-quality |
| bin.2.105 | *L. cremoris* | 67.92, 0.8 | 100 | 1 | 1713729 | 1713729 | Medium-quality |
| bin.s.4 | *L. cremoris* | 100, 0.08 | 0 | 1 | 2379957 | 2379957 | High-quality |
| bin.s.3 | *L. cremoris* | 100, 0.24 | 0 | 1 | 2412289 | 2412289 | High-quality |
| bin.s.2 | *L. cremoris* | 99.99, 0.43 | 0 | 1 | 2411749 | 2411749 | High-quality |
| bin.2.103 | *L. falkenbergense* | 61.32, 0.21 | 0 | 1 | 1233481 | 1233481 | Medium-quality |
| bin.3.28 | *L. falkenbergense* | 62.72, 8.26 | 100 | 4 | 772407 | 1456060 | Medium-quality |
| bin.s.25 | *L. falkenbergense* | 99.82, 0.3 | 0 | 1 | 1882740 | 1882740 | High-quality |
| bin.s.21 | *L. falkenbergense* | 99.88, 0.22 | 0 | 1 | 1972902 | 1972902 | High-quality |
| bin.s.22 | *L. falkenbergense* | 99.88, 0.22 | 0 | 1 | 1972903 | 1972903 | High-quality |
| bin.2.26 | *L. lactis* | 81.16, 0.23 | 0 | 1 | 1908490 | 1908490 | Medium-quality |
| bin.s.5 | *L. lactis* | 98.87, 3.4 | 78.95 | 1 | 2540971 | 2540971 | High-quality |
| bin.3.105 | *L. laudensis* | 54.51, 0.89 | 0 | 3 | 595058 | 1281166 | Medium-quality |
| bin.3.102 | *L. laudensis* | 60.52, 0.09 | 0 | 2 | 753661 | 1322493 | Medium-quality |
| bin.2.73 | *L. laudensis* | 66.62, 0.21 | 0 | 1 | 1454674 | 1454674 | Medium-quality |
| bin.2.88 | *L. laudensis* | 70.77, 0.05 | 0 | 1 | 1595247 | 1595247 | Medium-quality |
| bin.2.9 | *L. laudensis* | 73.75, 0.32 | 0 | 1 | 1663044 | 1663044 | Medium-quality |
| bin.3.160 | *L. laudensis* | 75.47, 0.57 | 0 | 2 | 1015763 | 1566712 | Medium-quality |
| bin.s.7 | *L. laudensis* | 99.72, 1.18 | 0 | 1 | 2301720 | 2301720 | High-quality |
| bin.2.6 | *L. mesenteroides* | 81.71, 0.2 | 0 | 1 | 1515853 | 1515853 | Medium-quality |
| bin.s.24 | *L. mesenteroides* | 97.33, 0.27 | 0 | 1 | 1629765 | 1629765 | High-quality |
| bin.s.23 | *L. mesenteroides* | 97.45, 0.13 | 0 | 1 | 1629550 | 1629550 | High-quality |
| bin.1.77 | *L. mesenteroides* | 99.06, 3.73 | 92.31 | 2 | 1251071 | 1742053 | High-quality |
| bin.s.17 | *L. mesenteroides* | 99.85, 0.34 | 0 | 1 | 1930162 | 1930162 | High-quality |
| bin.s.18 | *L. mesenteroides* | 99.92, 0.36 | 0 | 1 | 1891954 | 1891954 | High-quality |

In parallel, we expanded upon previously reported collections of *L. cremoris, L. mesenteroides*, and *P. laudensis* obtained from Tistrup [12, 18]. The original isolations on LM17 and GMA focused on extensive colony sampling, whereas the expanded isolations tested additional media and incubation conditions with fewer isolates sampled per condition, targeting *L. lactis* and additional strain diversity among the rest of the community members. Recovery varied markedly among species: despite its substantial abundance in the metagenome, *L. lactis* was only recovered with selective plating at 20 °C with 3 % NaCl, whereas *L. falkenbergense* remained unrecovered (Sup. Fig. S1). Newly obtained isolates and existing draft genomes were (re-)assembled and the collection was dereplicated using genomic and epigenomic criteria to retain non-redundant representatives for downstream analyses (See Methods). This resulted in 111 non-redundant isolates out of 199 total isolations, comprising 55 *L. cremoris*, 5 *L. lactis*, 36 *P. laudensis*, and 16 *L. mesenteroides* (Fig. 1c; Suppl. Table S1).

We next investigated whether cultivation and metagenomics recovered the same below-species population structure. Minimum within-species ANI values in *L. cremoris* (98.72 %), *L. mesenteroides* (98.72 %), and *P. laudensis* (99.06 %) indicated appreciable genomic divergence among coexisting strains, while in *L. lactis* (99.79 %) it was more limited (Sup. Fig. S2). Core-genome phylogenetic analysis and fast hierarchical Bayesian analysis of population structure of the isolates resolved 14 within-sample population clusters (hereof core-genome lineages): four *L. cremoris*, three *L. mesenteroides*, five *P. laudensis*, and two *L. lactis* lineages. Incorporation of high-quality MAGs without strain heterogeneity expanded this structure to 16 lineages across the four species. Only five core-genome lineages had representatives from both isolates and MAGs, whereas two were exclusively recovered by metagenomics and nine by cultivation (Sup. Fig. S3). Thus, even at the core-genome lineage level, the two approaches captured different but complementary fractions of the diversity.

To identify whether the partial overlap observed in core-genome linages extended to genetic content, we compared species-level isolate gene pools against curated, chromosomal-only metagenome gene catalogues. Across species, metagenomics recovered substantial additional gene-family diversity compared to isolate WGS, with the largest difference observed in *L. cremoris* (Fig. 1d), suggesting the largest underlying genetic diversity not captured by cultivation in this species, at least at chromosome level. Functional annotation of the metagenome-only genes assigned 65.0-81.6 % of them to clusters of orthologous groups (COGs) categories, revealing substantial representation of functions related to transport and metabolism, mobilome, and cell wall and envelope biogenesis, The latter included genes associated with surface polysaccharide biosynthesis, such as cell wall polysaccharides (CWPS) and exopolysaccharides (EPS) (Fig. 1c). Altogether, these results indicated that the incomplete recovery of community diversity extended to accessory functions associated with gene flux, metabolism and cell-surface variation, prompting the examination of both phage-interaction loci and the mobilome in greater detail.

### Cell wall polysaccharide diversity reveals substantial cultivation bias

CWPS constitute the principal phage receptors for prevalent phages infecting dairy *Lactococcus,* including *Skunavirus*, and their biosynthetic machinery is encoded within the so-called *cwps* locus [19, 20]. Because no established *cwps* identification nor classification framework exists for *Leuconostoc*, we excluded identified putative cell-surface polysaccharide-encoding loci of *L. mesenteroides* from the results, even though substantial diversity was noted in evident polysaccharide biosynthesis loci (data not shown). In *Lactococcus* and *Pseudolactococcus*, we identified *cwps* clusters and categorized them according to previously established genotype classifications [12, 19] (Sup. Fig. S4).

Two major CWPS genotypes were identified in *L. cremoris* isolates (A and C), both being subdivided into C_1_ and C_2_ and A_1_ and an unclassified type A genotype (AU) (Figure 2a). In *P. laudensis, cwps* clustered into two major genotypes (F and G), with F subdivided into subgroups (F-a to F-d) depending on the position of transposase-associated insertions within the loci. The subgroups were retained due to potential alteration of CWPS structure and thus phage adsorption properties. *L. lactis* strains contained exclusively B_1_ genotypes. Despite substantial locus-level variation, *cwps* genotype was largely associated with core-genome population structure (Fig. 2b) and therefore captured broad population structure.

**Figure 2.**
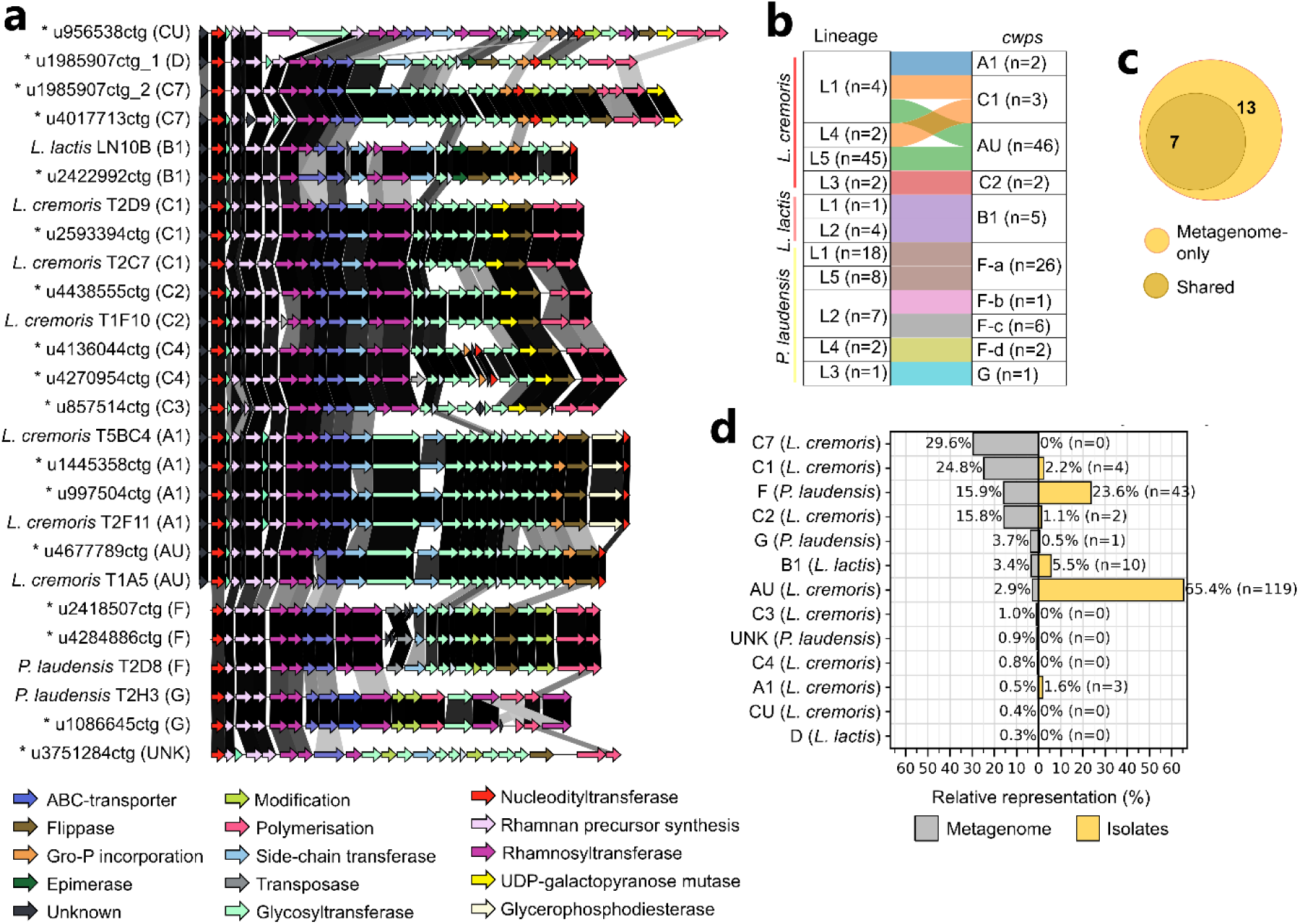
Diversity and recovery of *cwps* genotypes in Tistrup. **a** Gene-level comparison of representative CWPS biosynthesis loci from isolate genomes and metagenomic contigs (asterisks). Genotype designations are shown in parentheses. Arrows indicate gene orientation and are coloured by functional annotation, and shaded links connect homologous regions between loci. **b** Relationships between core-genome lineages and *cwps* genotypes in the non-redundant isolate collection. Links represent observed lineage–genotype associations and are not weighted by isolate counts; values in parentheses indicate the number of isolates per category. **c** Overlap of *cwps* genotypes recovered from isolates and metagenomic contigs, with numbers indicating genotype counts. **d** Relative representation of *cwps* genotypes in the metagenome and isolate collection before dereplication. Metagenomic percentages represent mean locus depth normalized by the summed mean depth across reference genotypes. Isolate percentages are calculated from 182 *cwps*-typed isolates pooled across cultivation conditions, with counts shown in parentheses. Sampling effort varied among cultivation conditions. The F-a–F-d subtypes distinguished in b are pooled as F in a and d.

We next assessed how completely this diversity was represented by cultivation. The metagenome analysis recovered all seven *cwps* genotypes observed among isolates and revealed seven additional variants not captured by cultivation (Figure 2ac). Specifically, five *cwps* genotypes (C_7_, C_4_, C_3_, and a novel C subtype) were identified in contigs taxonomically assigned to *L. cremoris*, one genotype (D) to *L. lactis*, and a novel genotype in contigs assigned to *P. laudensis* (UNK) (Figure 2a). Read-based quantification of metagenomic *cwps* depth revealed an even stronger discrepancy between community representation and isolate recovery (Fig. 2d). Despite C_7_ being the most highly represented genotype in the metagenome (29.6 % of the total), this was absent from the isolate collection. Similarly, C_1_ and C_2_ were abundant in the metagenome (24.8 % and 15.8 %, respectively) but represented by only four and two isolates. Conversely, AU accounted for only 2.9% of *cwps* in the metagenome but was originally represented by 119 isolates (46 different after dereplication), representing 65.4 % of the *Lactococcus/Pseudolactococcus* isolations (Figure 2d). Altogether, these results identify CWPS as a population-structured but incompletely sampled layer of below-species diversity. The metagenome revealed the multiple CWPS backgrounds absent from the isolate set, and the marked discordance between metagenomic abundance and isolate recovery exposed important cultivation biases, including complete failure to recover the dominant CWPS genotype.

### Additional phage interaction layers provide extensive diversity beyond core-genome population structure

While *cwps* diversity broadly tracked core-genome population structure, we next investigated whether other genomic features involved in phage–host interactions followed the same pattern or revealed additional variation among closely related strains.

EPS, whose biosynthesis is encoded in the *eps* cluster, and which can influence phage adsorption by masking receptors or acting as adsorption substrates [21, 22], were variably present across the isolate collection and showed little correspondence with core-genome population structure. EPS-encoding clusters occurred in most *L. lactis* and *P. laudensis* isolates but only a subset of *L. cremoris*, resolving into 12 distinct genotypes (Fig. 3a). The metagenomic analysis recovered most isolate-associated genotypes while identifying 16 additional variants across *Lactococcus* and *Pseudolactococcus* contigs (Fig. 3c; Sup. Fig. S5), including shared genotypes in *L. lactis* and *L. cremoris* contigs, in agreement with interspecies HGT of the loci.

**Figure 3.**
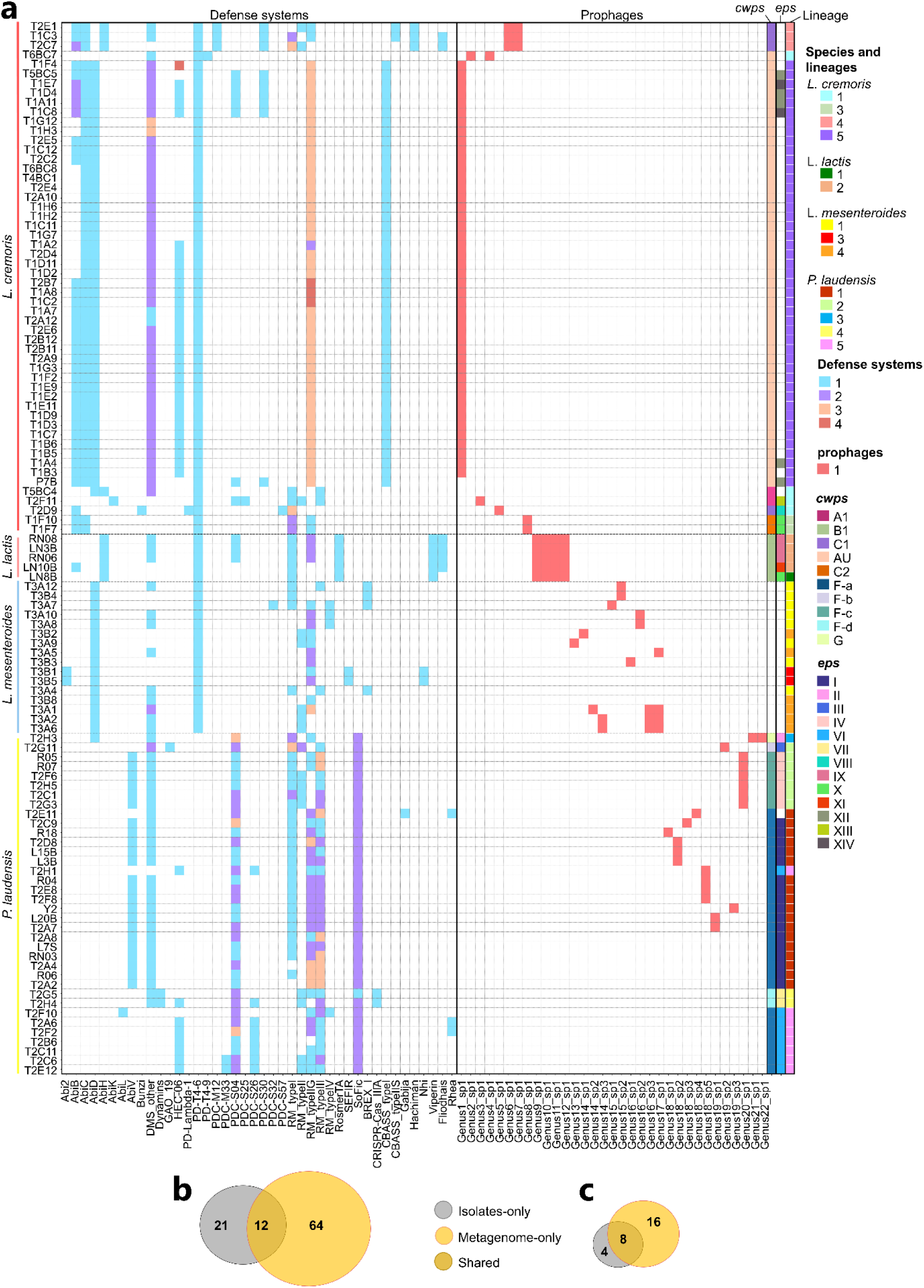
Diversity and recovery of phage-interaction determinants in Tistrup. **a** Distribution of predicted antiphage defence systems, intact prophages and surface polysaccharide genotypes across non-redundant isolates, grouped by species, and clustered by similarity in defence systems counts. Rows represent isolates; columns show defence-system types (left) and prophage species-level clusters labelled by genus and species cluster identifiers (middle). Defence-system colours indicate copy number (1–4), whereas red cells indicate prophage presence. White cells in these matrices indicate no detection. Annotation tracks on the right show *cwps* and *eps* genotypes and core-genome lineage assignments. CWPS and EPS-encoding clusters were not analysed in *L. mesenteroides*. b Overlap between isolate and metagenomic datasets for prophage species-level clusters. c Overlap between isolate and metagenomic datasets for *eps* genotypes. Numbers indicate clusters or genotypes detected exclusively in isolates, exclusively in the metagenome, or shared in both datasets.

Prophage repertoires provided another highly variable component of the isolate genomes. Intact prophages detected across the collection represented 22 viral genera and 33 viral species, with most isolates carrying at least one prophage and *L. lactis* carrying the highest burden with four prophages per strain. Prophage composition at both genus and species level were often core-genome lineage-independent (Fig. 3a). Comparison to the metagenome revealed only a 12 species overlap in prophage repertoires, while uncovering a markedly larger diversity (64 additional species) than captured by the isolate collection (Fig. 3b). Proteome-based phylogenetic analysis of the prophages showed structuring according to host genus, while prophages associated with *L. lactis* and *L. cremoris* were intertwined (Sup. Fig. S6), suggesting evolutionary relatedness across prophages from the two *Lactococcus* species.

Antiphage defence repertoires were likewise heterogeneous across the isolate collection, even among closely related isolates, providing another layer of diversity not captured by core-genome structure alone. In total, we identified 34 established defence systems and eight additional phage-defence candidates (PDCs), with substantial variation both within and between species (Fig. 3a). *L. cremoris* encoded the highest number of systems per strain on average (10), followed by *P. laudensis* (8.25), *L. lactis* (7.8) and *L. mesenteroides* (5.25). Restriction–modification (RM) and other DNA methylation-associated systems (DMS) dominated the defensome, although their composition differed among species, while abortive infection (Abi) systems were particularly frequent among the lactococci. All the major defence-system classes observed in isolates were also recovered in the metagenome, which furthermore revealed systems absent from the cultivated strains. Additional Abi systems, including AbiE, AbiJ and AbiN, were detected only metagenomic contigs assigned to either the lactococci or *P. laudensis*.

RM and other methylation-based systems generate methylated DNA motifs in the genome of their hosts, providing an additional measurable epigenetic layer of diversity. DNA methylation analysis was performed for 56 isolates, including 11 *L. cremoris*, 34 *P. laudensis*, five *L. lactis* and six *L. mesenteroides*. Across these isolates, we detected 73 methylated motifs, no methylated motif was universally conserved across all isolates within any species, and core-genome lineage-specific motifs were likewise rare. The latter were observed only for GGCT6mA and TC6mANNNNNNNTGC within *L. cremoris* lineage “5”, GTAAN6mAC within *L. mesenteroides* lineage “4”, and CTAY6mAB in *L. lactis* lineage “2”. Only the motif TC6mANNNNNNNTGC was shared across species, concretely between some *L. lactis* and *L. cremoris* strains (Sup. Fig. S7). Methylome composition was instead highly strain-specific: 38 of 56 isolates exhibited a unique motif repertoire (Sup. Fig. S7). Comparison with the metagenomic methylome exhibited limited overlap under the applied detection thresholds, with only 20 motifs detected in both datasets, 77 exclusively in the metagenome, and 53 only among isolates (Fig. 4a). The large number of dataset-specific motifs further indicated that neither approach captured the complete methylation repertoire of the community.

**Figure 4.**
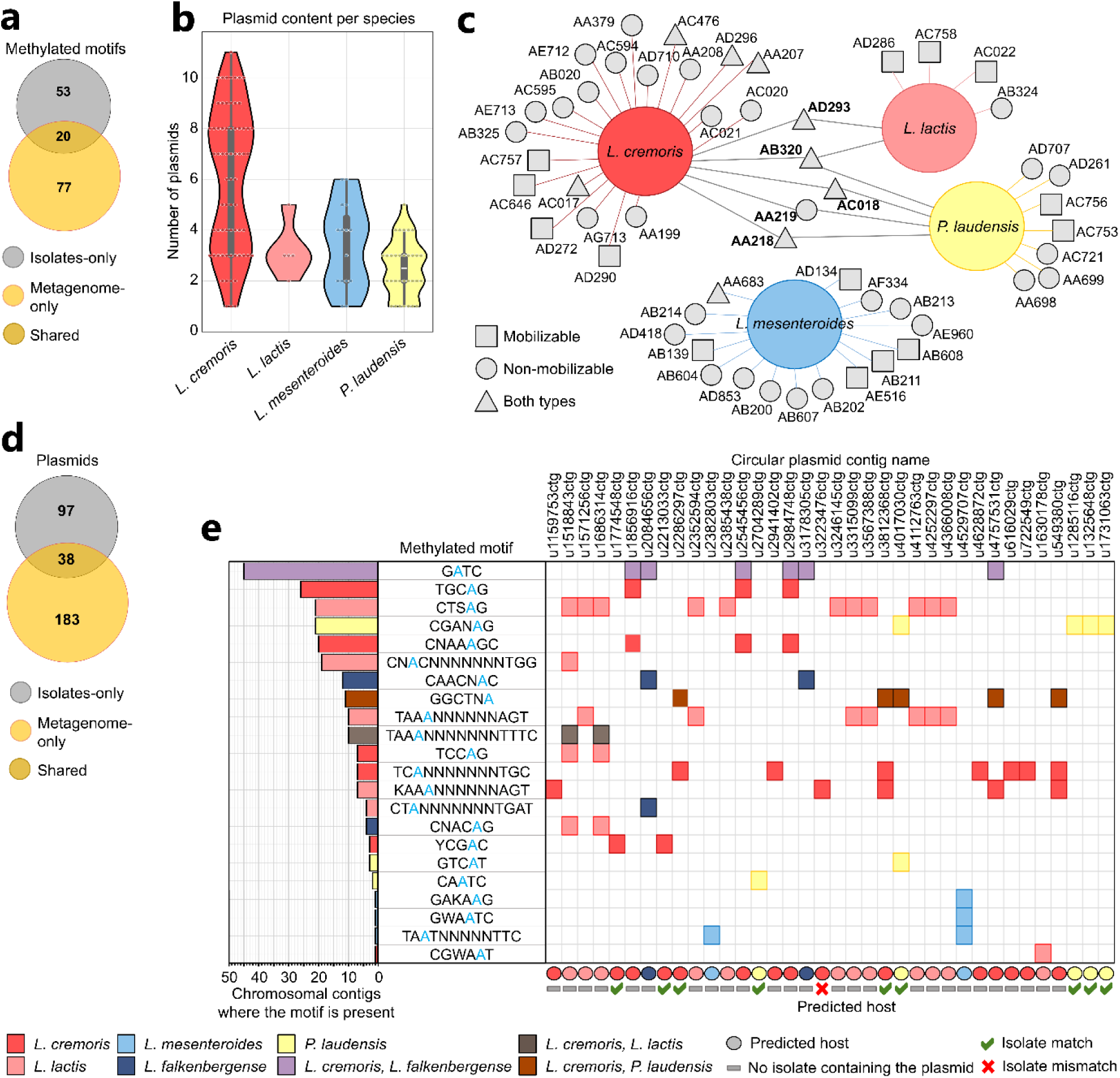
Plasmid diversity, methylation signatures and inferred host associations. **a** Overlap in methylated motifs detected in isolate and metagenomic datasets. **b** Distribution of plasmid counts per species. Dots represent individual isolates, embedded boxplots indicate medians and interquartile ranges, and violin widths represent density normalized within species and do not reflect sample size. **c** Distribution of MOB-suite primary plasmid groups across species in the isolate collection. Links connect species to the plasmid groups recovered from their isolates; bold labels identify groups shared across species. Node shapes indicate predicted mobility of group members. **d** Overlap between distinct plasmids recovered from isolates and dereplicated circular plasmids reconstructed from the metagenome. **e** Methylation-based host assignment for 36 circular metagenomic plasmids. Horizontal bars indicate the number of chromosomal contigs on which each methylated motif was detected, while the matrix shows detection of these motifs on individual plasmid contigs. Colours indicate the species or combinations of species associated with each motif on chromosomal contigs, and blue letters mark modified bases (6mA). White cells indicate no detected methylation for the corresponding motif. Coloured circles below the matrix indicate predicted host species. Green ticks and a red cross indicate agreement and disagreement, respectively, with host associations observed in isolates, while grey dashes indicate plasmids not recovered from the isolate collection.

Overall, EPS, prophages, defence systems, and methylation repertoires revealed extensive variation beyond that captured by core-genome population structure or CWPS genotype, providing multiple complementary dimensions of below-species diversity.

### A diverse plasmidome carries key fermentation genes and metagenomic methylation signatures support host assignment

Beyond variation in phage-interaction determinants and methylation profiles, the mobile accessory genome represented another major layer of below-species diversity within the community. Plasmids were abundant across the Tistrup isolates and varied substantially among species, with *L. cremoris* isolates carrying an average of approximately six plasmids per strain, compared with 3.4 in *L. mesenteroides*, 3.2 in *L. lactis* and 2.5 in *P. laudensis* (Fig. 4b). Following dereplication (see Methods), the isolate collection contained 135 distinct plasmids ranging 3,084-71,694 bp, distributed across 53 MOB-suite [23] primary groups or “cluster IDs” (Sup. Fig. S8), revealing extensive plasmid diversity in the community. In general, plasmids were a major contributor to strain differentiation across the dataset (Sup. Fig. S8). For instance, within *L. cremoris* lineage “5”, strains exhibited near-identical chromosomal sequences (ANI 99.95-100 %) and these were primarily differentiated by their plasmid complement.

Interspecies sharing of plasmid groups was uncommon, with only five groups detected across multiple hosts (Fig. 4c). Three of the groups (AA218, AA219 and AC018) were observed in both *P. laudensis* and *L. cremoris* but showed limited genetic homology beyond replication-associated genes, transposases, and small hypothetical proteins. In contrast, group AD293 was shared between *L. lactis* and *L. cremoris* and contained several near-identical loci, including *prtP*, encoding the cell-envelope proteinase involved in casein degradation [24], and a putative transposon carrying copper-export genes. Group AB320 exhibited the broadest host range (Fig. 4c). *L. lactis* and a subset of *L. cremoris* harboured near-identical AB320 plasmids (with minor rearrangements), containing the *lac* operon for lactose utilization and *opp* transporters for oligopeptide uptake, consistent with HGT between these taxa. *P. laudensis* AB320 plasmids were more divergent, sharing primarily replication genes and the *lac* operon with the lactococcal plasmids. Dairy-relevant functions were not restricted to these shared plasmid groups but were also distributed among species-specific plasmids. For example, *prtP* was also detected in *L. cremoris* AC021 and AC757 plasmid groups, EPS-associated loci in *L. cremoris* AC757 and *L. lactis* AC758 groups, and lactose transport and metabolism modules in *L. mesenteroides* AD134, AB604, AB202, AE516 and in *P. laudensis* AA219 and AD297 groups.

To compare plasmid recovery from isolate sequencing versus metagenome assembly, we identified plasmid contigs in the metagenome, yielding 1,107 plasmid-classified contigs out of 2,691 total contigs, including 253 circular plasmids. After dereplication, we distinguished 221 distinct circular plasmids representing 58 MOB-suite groups, of which 41 were also represented among the isolates. Despite this overlap at the plasmid-group level, concordance at the individual plasmid level was considerably lower: only 38 of the 135 isolate plasmids were recovered as circular metagenomic plasmids (Sup. Fig. S9), leaving 97 represented only among isolates and 183 detected only in the metagenome (Fig. 4d). Therefore, as observed for chromosomal population diversity, cultivation and metagenomics recovered strongly complementary fractions of the community plasmidome.

An added challenge when assessing the complementarity of the datasets is that the cellular hosts of metagenome-derived plasmids cannot be determined directly from their assembled sequences.

Because DNA methylated motifs are shared across intracellular replicons [25], we leveraged species-resolved methylome “fingerprints” to infer the hosts of metagenome plasmids. Following taxonomic annotation of contigs assigned as chromosomal, we identified methylated motifs only occurring in subsets of contigs corresponding to a particular species, including sets unique to *L. mesenteroides* (17 motifs), *P. laudensis* (20), *L. lactis* (26), *L. cremoris* (22) and *L. falkenbergense* (4), alongside a smaller subset shared across species (Sup. Fig. S10).

Methylation was detectable on 36 of the 221 circular plasmid contigs, enabling host prediction based on the presence of species-associated signature motifs (Fig. 4e). Using this approach, we assigned 16 plasmids to *L. cremoris*, 11 to *L. lactis*, 5 to *P. laudensis,* 2 to *L. mesenteroides,* and 2 to *L. falkenbergense*. Ten of these plasmids were also recovered among isolates, providing an opportunity for independent validation, of which nine supported the methylome-based host assignment (Fig. 4e), whereas one plasmid (contig u3223476ctg; MOB-suite group AC758) was predicted to originate from *L. cremoris* but was recovered from *L. lactis* isolates. This discordance is consistent with interspecies transfer of the plasmid, particularly since the plasmid encodes an apparent mobilization module (comprising an oriT, relaxase, type IV coupling protein, and auxiliary relaxosome-associated genes). Additional isolate sampling would therefore be required to resolve its host distribution. Even though host associations were concordant for most motifs shared across datasets, three motifs (G6mATGG, TC6mANNNNNNNTGC and TTC6mANNNNNNRTC) showed broader host distributions in one dataset than the other, increasing the number of plausible motif–species associations.

Beyond species-level host assignment, methylation signatures revealed plasmid–methylome combinations absent from the isolate collection. Among the nine metagenomic plasmids with isolate-supported host assignments, two *P. laudensis* AC721 plasmids (u4017030ctg and u1285116ctg) contained the CGAN6mAG motif. This motif occurred in a subset of isolates, yet none of them harboured the corresponding plasmids. Similarly, the u1774548ctg and u2213033ctg *L. cremoris* plasmids contained the YCG6mAC motif, which was not detected in any analyzed isolate. Thus, these examples represented plasmid–methylome combinations not captured among the sequenced isolates, suggesting their occurrence in unsampled community members.

Overall, these results reveal an extensive plasmidome involved in the dissemination of dairy-relevant functions, with cultivation and metagenomics capturing complementary diversity and methylation enabling host assignment at species level and detection of unsampled plasmid–methylome combinations.

### Genomic and epigenomic markers provide complementary strain-level resolution

Across the isolate collection, each of the genomic and epigenomic features examined above varied among otherwise closely related strains, suggesting that they capture distinct layers of below-species diversity. We therefore asked how much discriminatory resolution each marker provided individually and how much additional resolution was gained when markers were combined. CWPS, EPS and prophage repertoires generally resolved fewer profiles, whereas plasmid-group, defence system and methylome repertoires provided higher discrimination, although their relative discriminatory power differed among species. Combining the markers, 103 out of 111 isolates possessed uniquely resolved strain-level profiles (Sup. Data, not available in the preprint version), while the rest were distinguished by genome length and ANI criteria (Methods). Pairwise combinations frequently resolved additional profiles beyond either marker alone (Fig. 5), demonstrating that these features capture complementary dimensions of within-species diversity. Among isolates with complete data across all studied markers (n=55), methylomes provided the highest resolution in *L. mesenteroides* and *P. laudensis*, whereas plasmid-group repertoires were the most discriminatory in *L. cremoris* and *L. lactis* (Fig. 5), and the combination of the markers distinguished every isolate. Therefore, population diversity in the Tistrup community is distributed across multiple complementary genomic and epigenomic layers that together provide greater strain-level resolution than any marker alone.

**Figure 5.**
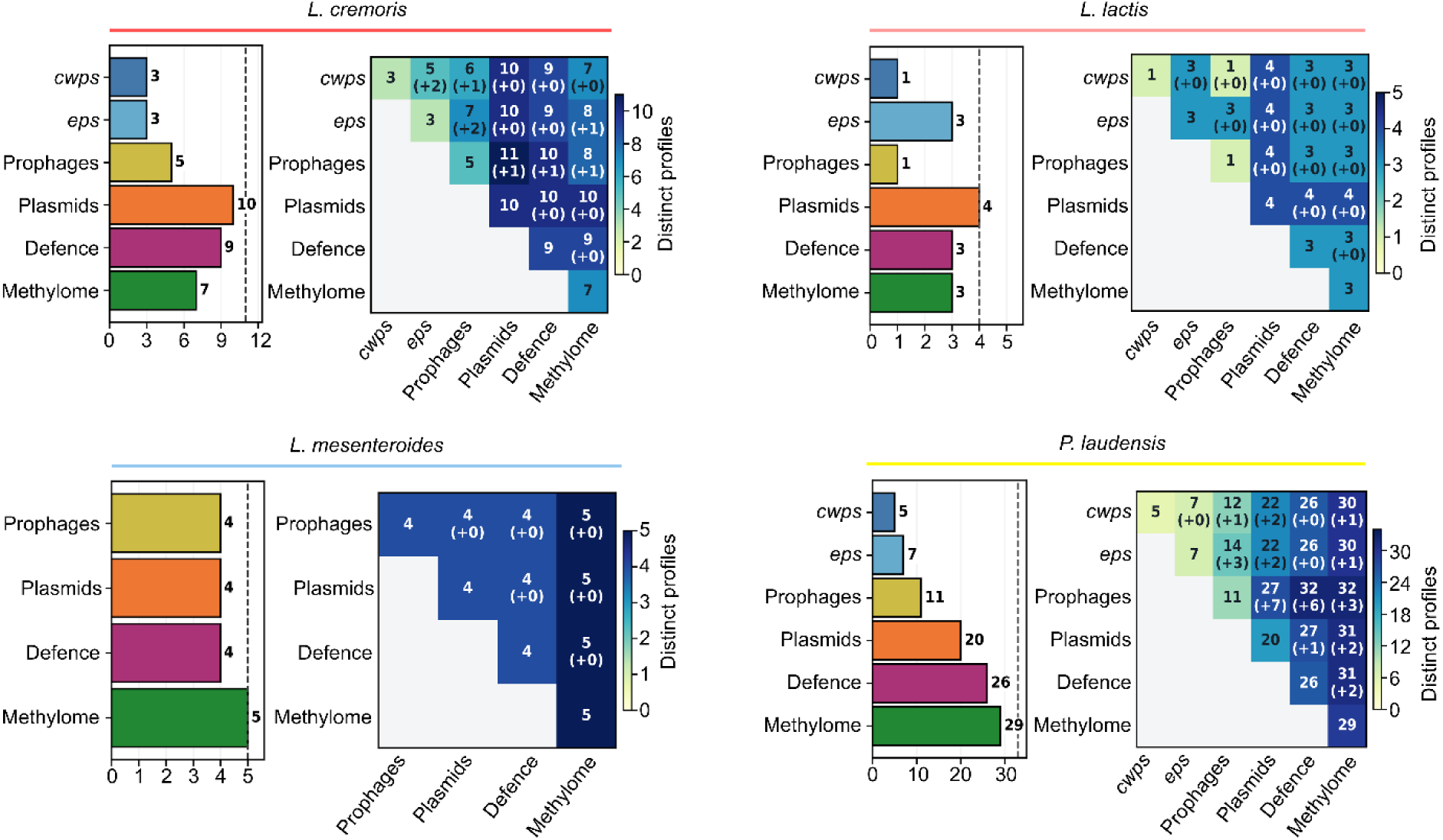
Complementary resolution of strain-level diversity by genomic and epigenomic markers. For each species, (left) Numbers of distinct profiles resolved by individual marker categories. Dashed lines indicate the number of profiles resolved by combining all displayed categories. (right) Pairwise combinations of marker categories. Cell colours and values indicate the number of distinct combined profiles, while values in parentheses indicate the increase relative to the more discriminatory individual marker. Comparisons within each species use the same isolates with complete data for all displayed categories: *P. laudensis*, n = 34; *L. lactis*, n = 5; *L. cremoris*, n = 11; and *L. mesenteroides*, n = 5. *eps* and *cwps* were not analysed in *L. mesenteroides* and are omitted. Confirmed absence constitutes a valid profile, whereas missing data are excluded. Colour scales are specific to each species.

## Discussion

Despite major advances in long-read sequencing and genome-resolved metagenomics, comprehensive reconstruction of strain-level diversity and mobile genetic repertoires remains challenging. Most deeply studied microbiomes, including the human gut and soil, are taxonomically rich and can yield hundreds of distinct MAGs, but their low cultivability hinders complementary isolate-based interrogation and validation of fine-scale population diversity below the species level [26]. Here, we examined the opposite community paradigm using a mesophilic dairy starter culture dominated by a few, highly culturable species as a model to resolve the extent and structure of below-species level diversity in an ecosystem shaped by persistent phage pressure. [14, 20]. Our results show that low species richness in Tistrup conceals extensive combinatorial variability in genomic and epigenomic features linked to fermentation traits and phage-host interactions among closely related strains.

Intensive cultivation and deep long-read metagenomics captured complementary but incomplete fractions of Tistrup diversity. Cultivation favoured populations able to grow under the conditions tested, whereas high sequence similarity among coexisting strains constrained metagenomic reconstruction. Biases extended beyond rare community members: *L. lactis* represented ∼20% of the read-based community signal but accounted for limited isolate diversity, while *L. falkenbergense* was not recovered despite a read-based abundance approximately tenfold lower than that of *L. mesenteroides*. Below the species level, different cultivation conditions recovered distinct core-genome lineages, with limited overlap between the lineages represented by isolates and MAGs. CWPS biosynthesis genotypes broadly tracked core-genome structure, yet their representation differed markedly between datasets to the point in which the dominant *L. cremoris cwps* genotype in the metagenome, C_7_, accounted for nearly 30% of the CWPS signal but was completely absent from the isolate collection. Conversely, genotypes prevalent among isolates contributed only minor fractions of that signal, in agreement with previous observations in dairy starters [27]. Thus, extensive isolate collections can omit abundant genotypes, while deep metagenomics can detect diversity without resolving the corresponding strain genomes. A further dimension of heterogeneity emerged from the combinations of genomic and epigenomic features within closely related bacterial backgrounds.

Cultivation remained essential for linking CWPS and EPS genotypes, plasmid and prophage repertoires, defence systems and methylation profiles within individual strains, revealing distinct combinations among closely related isolates. These features can be genetically and functionally interconnected: MGEs carry most defence systems and all EPS biosynthesis clusters described to date in *Lactococcus* and *Pseudolactococcus* [9, Soto-Serrano et al., in press], surface polysaccharides and defence systems can influence susceptibility to specific phages [19, 28, 29], and RM systems dictate methylation profiles. Despite these connections, their combinations, potentially shaped by HGT and persistent phage pressure, provided complementary discriminatory power. The most discriminatory marker varied among species, and pairwise combinations frequently resolved additional profiles, whereas integration of all available markers distinguished every non-redundant isolate with complete data i.e., those containing all studied features. These results demonstrate that the combinatorial variability of genomic and epigenomic markers associated with phage resistance and key fermentation traits provides a biologically relevant basis for strain-level discrimination in Tistrup. Accordingly, we propose combining these markers to resolve strain diversity in dairy starter communities more broadly.

Long-read metagenomics extended this view beyond cultivated strains, recovering additional polysaccharide genotypes, prophages, defence systems and plasmids. The extensive diversity recovered across these features, together with the limited overlap between datasets, exposed a vast population heterogeneity that remained hidden to either approach alone. Moreover, even where genetic content overlapped, metagenomics revealed additional host contexts, showing that recovering shared features does not ensure complete representation of the populations carrying them. Although metagenomic methylation analysis was sensitive to coverage, calling thresholds, and composite signals from coexisting strains [2], with detectable motifs in only 16.3 % of the circular plasmids, species-level host assignments were corroborated by isolate genomes in nine of the ten testable cases, supporting their reliability. Methylation signatures also revealed plasmid–methylome combinations absent from the isolate collection, suggesting additional host backgrounds for an already represented plasmid group, exemplified by *P. laudensis* AC721 plasmids.

Low species richness and limited within-species diversity at sequence level, as reported for thermophilic starter cultures [30], are compatible with extensive variation in defence repertoires among closely related cheese-associated bacteria [31]. The findings presented here extend this perspective by resolving combinations across multiple genomic and epigenomic features within individual strain backgrounds. Population complexity in Tistrup therefore encompasses both variation in individual features and the configurations in which they occur, adding a dimension of heterogeneity that sequence relatedness does not capture. Even this expanded view likely represents only a lower bound on the population complexity of Tistrup, given the incomplete characterization of several community fractions. The absence of *L. falkenbergense* isolates impeded strain-level characterization of its accessory genome and methylome markers, while long-read taxonomic profiling suggested the existence of additional low-abundance taxa within *Lactococcus, Pseudolactococcus* and *Leuconostoc* that were represented by neither isolates nor resolved MAGs. Plasmid reconstruction was also incomplete: only 253 of 1,107 plasmid-classified contigs were circular, and many isolate plasmids were absent from the circular metagenomic assemblies and vice versa. Given the extensive heterogeneity within the populations examined most deeply in this study, these unresolved community fractions are most likely harbouring further variation in both individual features and their combinations.

In conclusion, this study illustrates how microbiomes with low species richness can conceal extensive genomic and epigenomic complexity that remains only partially resolved even by intensive cultivation and deep long-read metagenomics. The combinatorial variability of MGEs and phage-interaction determinants provides a plausible molecular basis for phage resilience in undefined starter cultures such as Tistrup, potentially limiting the fraction of the community susceptible to any one phage.

## Materials and methods

### Metagenomic DNA extraction

Metagenomic DNA extraction and all bacterial isolations used material from the same frozen batch of Tistrup starter culture (Arla Foods, Aarhus, Denmark). Metagenomic DNA was extracted using a modified phenol-chloroform method. Briefly, 10 ml of Tistrup starter culture liquid suspension stored at −60 °C were thawed on ice, mixed with 2 % citric acid, incubated at room temperature for 1 hour and centrifuged at 10,000 × g for 15 min. at 4 °C. The top layer was discarded, and the pellet was resuspended in 10 mL of PBS and centrifuged at 10,000 × g for 1 min to remove eventual floating fat. The sample was divided into 4 x 2 ml aliquots into clean Eppendorf tubes, mixed 1:1 with phenol, and incubated for 10 min in a fume hood with inversion every minute. After centrifugation (5,000 × g, 5 min), the aqueous phase was transferred to a new tube, extracted once with an equal volume of phenol–chloroform–isoamyl alcohol (25:24:1), and centrifuged again (5,000 × g, 5 min). The aqueous phase was recovered and DNA was precipitated by adding 2.5 volumes of ice-cold 96% ethanol followed by overnight incubation at −20 °C. DNA was pelleted by centrifugation (18,000 × g, 20 min), washed in 500 µl of 80 % ethanol, transferred to a 1.5 ml tube, and centrifuged again (18,000 × g, 10 min). After removing the ethanol, the pellet was air-dried and resuspended in 50 µl Milli-Q water. DNA concentration and purity were determined using a Qubit fluorometer (Thermo Fisher Scientific, Waltham, USA) according to the manufacturer’s instructions.

### Metagenomic DNA library preparation and sequencing

Extracted DNA was then subjected to library preparation using the Ligation Sequencing Kit V14 (SQK-LSK114; Oxford Nanopore Technologies, Oxford, UK) and sequenced using a PromethION 2 Solo platform (Oxford Nanopore Technologies, Oxford, UK). Basecalling was performed with Dorado v1.0.2 using the dna_r10.4.1_e8.2_400bps_sup@v5.0.0 basecalling model coupled with the dna_r10.4.1_e8.2_400bps_sup@v5.0.0_4mC_5mC@v3 and dna_r10.4.1_e8.2_400bps_sup@v5.0.0_6mA@v3 modified bases models

### Read-level taxonomic assignment and metagenome assembly

BAM files were converted to fastq format utilizing BEDtools v2.30.0 [32], and duplicated reads were removed with SeqKit v2.1.0 [33]. Nanofilt v2.8.0 [34] was used to filter the raw reads to a minimum quality of 18 and length of 1500 bases. Quality statistics of the reads were obtained with Nanoplot v 1.43.0 [34]. Read-level taxonomic classification was performed using Metabuli v1.1.1 [35] in precision mode (as defined by their authors: --seq-mode 3 --min-score 0.008) using a customized database generated by using Metabuli’s ‘updateDB’ function to add all complete genomes from the Lactobacillales family and all the draft and complete genomes obtained previously from the same starter culture to Metabuli’s prebuilt GTDB-based database [12, 18].

Filtered reads were assembled and binned with the mmlong2-lite pipeline v1.2.0 [26]. Briefly, Myloasm v0.2.0 [36] was used as an assembler, and Medaka polishing was disabled. Eukaryotic contigs were identified with Tiara v1.0.3 [37] and removed prior to binning. The remaining contigs were subjected to four successive binning rounds: (i) ensemble binning with MetaBAT2 v2.18 [38], SemiBin2 v2.2.0 [39] and Vamb v5.0.4 [40], followed by refinement with Binette v1.1.2 [41]; (ii) binning with COMEBin v1.0.4 [42]; (iii) a second ensemble round using the same tools and refinement procedure; and (iv) a final round using MetaBAT2 alone. At each round, bins passing CheckM2 v1.1.0 [43] completeness and contamination filters were retained (completeness ≥90% and contamination ≤5% in the first round, and completeness ≥50% and contamination ≤10% in subsequent rounds). Finally, taxonomic assignment of the bins was performed with GTDB-Tk v2.4.0 [44] against the Genome Taxonomy Database release 220.

### Bacterial isolation, DNA extraction, and genome sequencing

An initial set of 200 bacterial isolates was obtained from the same batch of frozen Tistrup starter culture (Arla Foods, Aarhus, Denmark) as described previously [12, 18]. Briefly, *L. cremoris* strains were isolated on a milk-based agar comprising skim milk supplemented with 0.2 % (w/v) tryptone, 50 mM MOPS (Sigma), and 0.003 % (w/v) bromocresol purple (Sigma-Aldrich, Saint Louis, USA), as well as on LM17 agar (M17 supplemented with 0.5 % (w/v) lactose). *P. laudensis* strains were isolated on LM17 agar. *L. mesenteroides* strains were isolated on MRS agar (HiMedia, Mumbai, India) supplemented with vancomycin 50 µg/mL. All the incubations were performed under anaerobic conditions using an AnaeroGen kit (Thermo Scientific, Waltham, USA). Colony-forming units (CFUs) were propagated thrice under the same conditions and subsequently incubated in their corresponding media in broth form for 48 h. The bacterial cells were then centrifuged at 4402g for 10 min utilizing a 5920 R centrifuge (Eppendorf, Hamburg, Germany), washed in 1ml 0.9% NaCl and centrifuged again at 12000 × g for 5min using a Micro Star 17R microfuge (VWR, Radnor, PA, USA). Subsequently, cell pellets were utilized for genomic DNA extraction using a Bead-Beat Micro AX Gravity kit following the instructions of the manufacturer (cat #106–100M1; A&A Biotechnology, Gdynia, Poland).

DNA sequencing library preparation was performed using Rapid Barcoding V14 (SQK-RBK114.96; Oxford Nanopore Technologies, Oxford, UK) and/or Native Barcoding V14 (SQK-NBD114.96; Oxford Nanopore Technologies, Oxford, UK) kits, following the instructions of the manufacturer and washing the final pooled libraries using Long Fragment Buffer for selection of fragments >3 kbp.

DNA sequencing was performed on a PromethION 2 Solo platform (Oxford Nanopore Technologies, Oxford, UK) and basecalled with Dorado v0.4.0 using the dna_r10.4.1_e8.2_260bps_hac@v3.5.2 basecalling model. Re-basecalling using newer models was not possible due to data being collected in the currently obsolete 4 kHz sampling frequency.

To broaden the isolate collection and target taxa within and beyond *L. cremoris*, *P. laudensis*, and *L. mesenteroides*, additional colony-forming units (CFUs) were recovered by plating the same batch of Tistrup starter culture on the following media and conditions: (i) MRS without vancomycin, incubated at 30 °C or 37 °C for 48 h; (ii) GLM17 (M17 supplemented with 0.5 % (w/v) lactose and 0.5 % (w/v) glucose) incubated at 42 °C for 48 h; (iii) Yeast Extract Lactate (YEL) agar, incubated anaerobically at 30 °C for 72 h using an AnaeroGen kit (Thermo Scientific, Waltham, USA) ; (iv) Reddy’s agar with or without 3 % (w/v) NaCl, incubated at 20 °C for 72 h; and (v) tryptic soy broth (TSB) agar, incubated at 37 °C for 48 h. Sampling effort varied among cultivation conditions (Sup. Fig. 1). DNA extraction was performed as described above, and the samples were subjected to library preparation using the Native Barcoding kit V14 (SQK-NBD114.96; Oxford Nanopore Technologies, Oxford, UK), following the instructions of the manufacturer. DNA sequencing was performed on a PromethION 2 Solo platform (Oxford Nanopore Technologies, Oxford, UK) and basecalled with Dorado v1.0.2 using the <u>dna_r10.4.1_e8.2_400bps_sup@v4.3.0</u> basecalling model coupled with the <u>dna_r10.4.1_e8.2_400bps_sup@v5.0.0_4mC_5mC@v3</u> and <u>dna_r10.4.1_e8.2_400bps_sup@v5.0.0_6mA@v3</u> modified bases models.

### Genome assembly and dereplication

The obtained BAM files were converted to fastq format utilizing Bedtools v2.30.0 [32], and duplicated reads were removed with SeqKit v2.1.0 [33]. Genome assembly was conducted using Hybracter v0.9.0 [45] with Flye [46] in “nano-hq” mode and Plassembler [47] using the plsdb_2023_11_03_v2 database. Minimum quality and read length were set at 13 and 1000 bp, respectively, and the subsample depth at 110×. Polishing with Medaka was omitted to avoid potential decrease in accuracy [3]. Genome assemblies exhibiting ≥1 non-circular contig were subjected to reassembly. For these, Nanofilt v2.8.0 [34] was used to filter the raw reads to a minimum quality of 8 and length of 1000 bp. Assembly was then performed using Trycycler v0.5.4 [48], after generating 15 read subsets and assemble them in groups of three utilizing Flye v 2.9.2 [46], Raven v1.8.1 [49], miniasm v0.3 + Minipolish v0.1.2 [50, 51], Hybracter v0.9.0 [45], and NextDenovo v2.5.0 + NextPolish v1.4.0 [52, 53].

To obtain a non-redundant dataset of strains for downstream analyses, isolates were dereplicated based on concordance across chromosomal and accessory-genome features. Two isolates were considered redundant if they (i) differed in chromosome or plasmidome length by ≤1000 bp, (ii) shared >99.99 % average nucleotide identity (ANI), and (iii) exhibited identical accessory profiles, defined as the same plasmid number and MOB-suite [23] primary cluster composition, prophage complement, defense-system repertoire (based on counts per defense system type), including DNA methylation motifs when available, as well as identical *cwps* and *eps* cluster genotypes.

### Phylogenetic analysis

For the construction of core-genome phylogenetic trees, pangenome analysis was performed at species level using Roary v3.13.0 [54] using a BLASTP identity of 90 % as a cutoff, using Bakta v1.9.4 [56] (v5.1 full database) gene annotations as an input. The resulting core alignment was subjected to analysis with Gubbins v3.4.3 [55] to identify and mask recombinant regions. The output recombination-filtered polymorphic alignment (SNP-only alignment) was used for phylogenetic reconstruction. Maximum-likelihood phylogenies were inferred with IQ-Tree v3.0.1 [56] using the GTR+G+ASC model. Branch support was assessed using 1,000 ultrafast bootstrap replicates and 1,000 SH-aLRT replicates, and trees were subsequently visualized and annotated in iTOL v6 [57]. To define within-sample lineages, the isolates were clustered using fast hierarchical Bayesian analysis of population structure using fastBAPS v1.0.8 [58] based on the recombination-filtered core-genome SNP alignment.

### Construction of metagenome-only species-specific gene pools

To construct species-associated chromosomal gene pools from metagenomic assemblies, contigs were taxonomically classified using Metabuli with the aforementioned custom reference database. Only contigs assigned to the focal species with a Metabuli v1.1.1 [35] classification score ≥0.7 and length ≥50 kb were retained for further analysis. To minimize contributions from MGEs, for which host in not certain, contigs predicted as plasmid- or virus-derived were excluded based on geNomad classification and only chromosomal contigs were used in the downstream analysis. The remaining contigs were independently validated by average nucleotide identity (ANI) against a collection of whole-genome sequences of the corresponding and similar species from both the isolate collection and the NCBI database using Skani v0.2.2 [59] in fast mode, requiring ANI ≥95 % and an aligned fraction ≥0.5 to the target species. Contigs also exhibiting exceeding the thresholds in closely related taxa were considered ambiguous and excluded from the corresponding species gene catalog to minimize cross-species misassignment. Contigs passing all filters were concatenated into a single multifasta file and annotated using Bakta v1.9.4 [60] in --meta mode to generate a metagenome-derived species gene catalog for downstream comparisons with isolate genomes.

Gene-family presence/absence matrices were inferred with Panaroo v1.5.1 [61] using a protein clustering identity threshold of 95 % (–c 0.95) and graph-based correction under clean-mode “moderate”, with paralogs merged (--merge_paralogs) and invalid gene models removed (--remove-invalid-genes). The resulting gene-family matrix was used to compare gene-family recovery between isolate WGS and metagenome-derived chromosomal gene pools. As a conservative measure to reduce false-positive spurious gene calls arising from fragmented metagenome assemblies and low-confidence gene prediction, hypothetical proteins shorter than 400 bp and only present in a single contig were excluded from the metagenome-derived gene catalog, thus prioritizing specificity in gene-family recovery estimates at the expense of underestimating the full metagenomic gene pool. Metagenome-only gene families were assigned to COGs using COGclassifier v2.0.0 (https://github.com/moshi4/COGclassifier) with the -e value set to 0.1.

### Surface polysaccharide-encoding biosynthetic gene clusters: *cwps*, and *eps*

The *cwps* loci encoding the necessary genes for CWPS biosynthesis was identified for isolates from the species in which this has been previously characterized i.e., *L. lactis, L. cremoris*, and *P. laudensis* [12, 19, 62]. Identification was performed as described elsewhere [Soto-Serrano et al., in press].

For identification in the metagenome assembly, all contigs >40 kb were identified in the assembly and annotated with Bakta v1.9.4 [60] in –meta mode and identification of *cwps* loci was performed as described elsewhere [Soto-Serrano et al., in press] and manually examined for completeness. Only clusters considered to encode a complete *cwps* cluster based on the presence of hallmark genes for both the locus and its surrounding regions [12] were utilized for downstream analyses.

Once *cwps* were identified in both metagenome and WGS isolate assemblies, nucleotide sequences were extracted with Samtools v 1.21 [63] and re-annotated with Bakta v1.9.4 [60] in –meta mode and resulting GenBank files were subsequently aligned and compared using Clinker [64] via the CAGECAT webserver, release 1.0 [65]. Annotation of the genes comprising the *cwps* loci was based on Bakta annotations, BLASTP alignments, and aided by transmembrane domains predictions using DeepTMHMM v1.0 [66].

For the classification of the *cwps* and *eps* loci into distinct genotypes, their extracted annotation files were analysed with Roary v3.13.0 [54] to create a binary presence/absence gene family matrix using a minimum percent identity for BLASTp of 50 % and without splitting paralogs. The presence/absence matrix was then used as an input for MeV suite v4.9.0 [67] and a two-way hierarchical clustering (HCL) was used to analyse and cluster the loci, using previously described genotypes as references [12, 19, 62, 68].

To establish the metagenomic abundance for each of the *cwps* genotypes, coverage values for the extracted *cwps* fasta files were generated using Samtools v1.22.1 [63] and Minimap2 v2.28 [69]. First, a multifasta file containing each of the genotypes as contigs was indexed (samtools faidx), and all reads were mapped to the file (minimap2 -ax map-ont). The resulting alignment file was then converted to BAM format (samtools view -b), sorted by coordinates (samtools sort), and indexed (samtools index). The sorted, indexed alignment was filtered to remove unmapped, secondary and supplementary records and keep only primary alignments (samtools view -F 0×904), to ensure that each read was only counted once in the coverage calculation. The filtered result was then used to compute per-genotype coverage statistics using samtools coverage.

#### Antiphage systems prediction and prophage prediction and classification

Antiphage systems were predicted utilizing PADLOC v2.0.0 [70] and DefenseFinder v2.0.1 [71], and where there were discrepancies in the presence of a specific system between the two tools, the positive result was selected. Furthermore, recently discovered antiphage systems in the lactococcal plasmidome [72] were detected using BLASTP [73] against the genes reported there; only matches with >90 % coverage and >99 % homology were accepted as positive.

Prophages from both isolates and metagenome were predicted with geNomad v1.8.1 [74] in conservative mode. Predicted sequences were both evaluated for completeness and treated for host contamination removal using CheckV v1.0.1 [75]. Host-free sequences classified as complete or high-quality (>90 % completeness) were further annotated with Pharokka v1.7.3 [76] and gene annotations were manually inspected. Only prophages encoding a whole set of structural genes were considered “intact” and kept for downstream analyses. Prophage sequences were clustered into genera and species using vClust v1.3.1 [77] by calculating the total ANI (tANI); species and genera were defined by ≥95 % and 70 % tANI, respectively. A proteome-based phylogenetic tree with species representatives was generated using ViPtree v4.0 [78] and edited with iTOL v6 [57].

#### DNA methylation analysis

DNA methylation was analysed for isolates sequenced under conditions enabling modified bases calling (i.e., Oxford Nanopore 5 kHz mode). In addition, methylome profiles previously generated for isolates from the same starter culture were incorporated to expand the isolate methylation dataset [79] [Li et al., under review]. Methylated motifs in the genome of the bacterial isolates were determined using Nanomotif v1.0.2 [2], following Nanomotif’s documentation and with default settings.

For metagenome-assembled contigs, methylated motifs were inferred with Nanomotif v1.0.2 using relaxed methylation detection thresholds (--methylation_threshold_low 0.2; -- methylation_threshold_high 0.35). A further reduction of --methylation_threshold_high to 0.25 did not materially change motif discovery and consequently 0.35 was retained. These relaxed classification thresholds were chosen because isolates from the same community have been shown to exhibit distinct methylomes [79], implying that strain-specific motifs could be present at modest read-level occupancy on shared contigs. Furthermore, the --min_motifs_bin was reduced to 15 to enable *de novo* motif discovery in smaller contigs. To compensate for these relaxed criteria, we enforced downstream stringency by modifying other discovery parameters as follows: -- minimum_kl_divergence 0.10, --min_motif_score 2, --threshold_valid_coverage 10.

#### Plasmid identification, typing, annotation, and host assignment

Plasmid contigs in the metagenomic assembly were predicted using geNomad v1.8.1 [74] in conservative mode and mob_recon v3.1.9 [23]. The predicted plasmids and those from the isolates were further typed with utilizing mob_typer v3.1.9 [23] to obtain information about mobilization genes as well as classifying the plasmids into MOB-suite clusters. Bakta v1.9.4 [60] was also used to perform gene annotations utilizing the v5.1 full database.

Only circular metagenomic plasmid contigs were retained for comparison with isolate plasmids. Plasmids were dereplicated separately within each dataset: those sharing both primary and secondary MOB-suite cluster identifiers and differing by <400 bp in length were considered redundant, with isolate plasmids additionally required to originate from the same host species. Plasmid overlap between datasets was defined by ANI ≥99.0% and an aligned fraction ≥80.0%. Pairwise comparisons and alignment visualizations were generated using FastANI v1.34 [80].

To assign metagenome plasmid contigs to hosts, we leveraged species-resolved methylome “fingerprints” derived from chromosomal contigs. First, chromosomal contigs for which DNA methylation was detected were taxonomically annotated using Metabuli v1.1.1 [35] and validated using manual BLASTN [73] searches. Subsequently, the methylomes of the chromosome contigs were scrutinized and species-specific motifs or combinations of motifs were ascertained independently of isolate-derived motifs. Plasmid-host links were inferred by comparing plasmid methylomes to the species fingerprints.

**The authors declare no competing interests**

---

**Supplementary Table S1.**
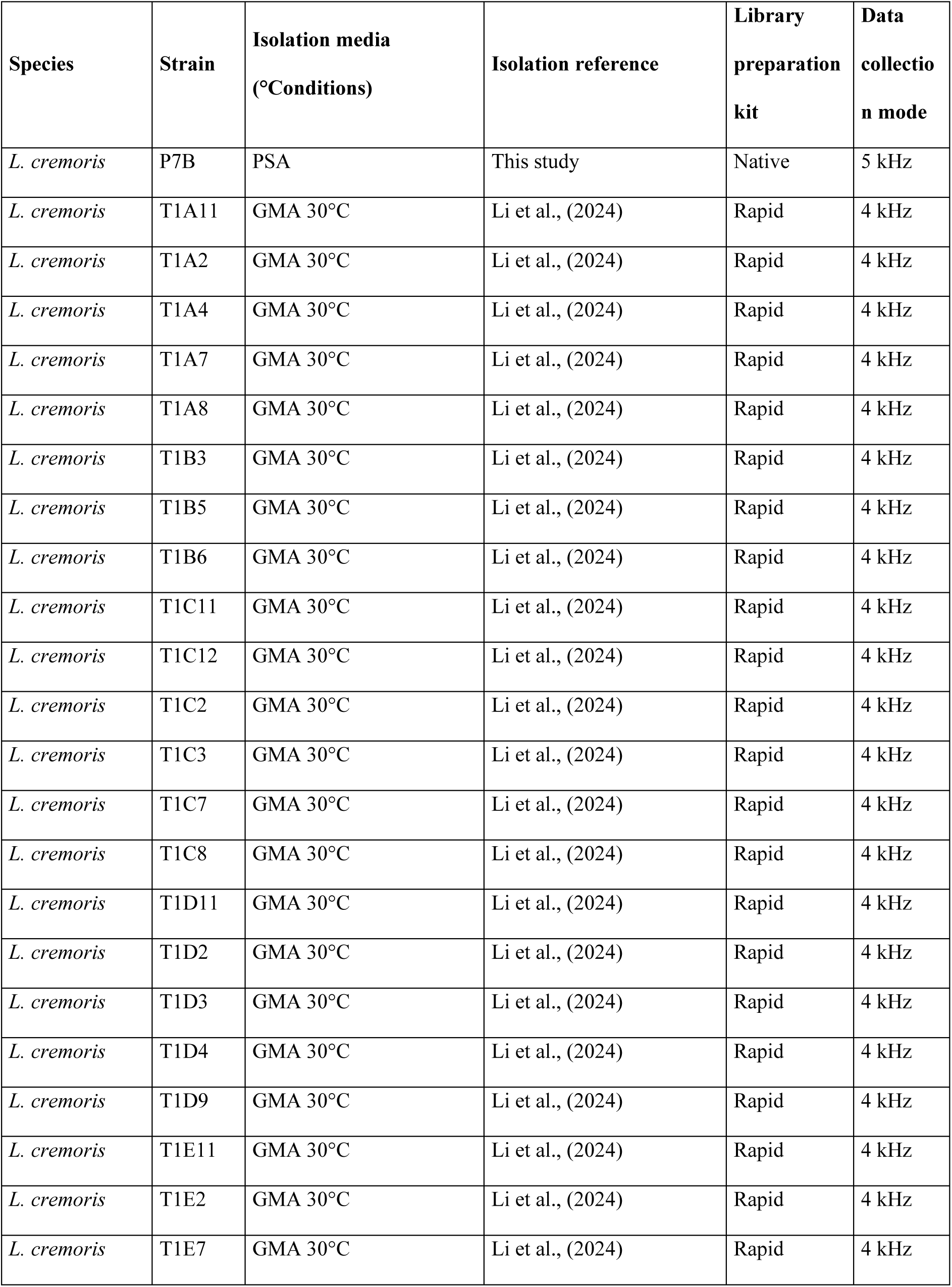
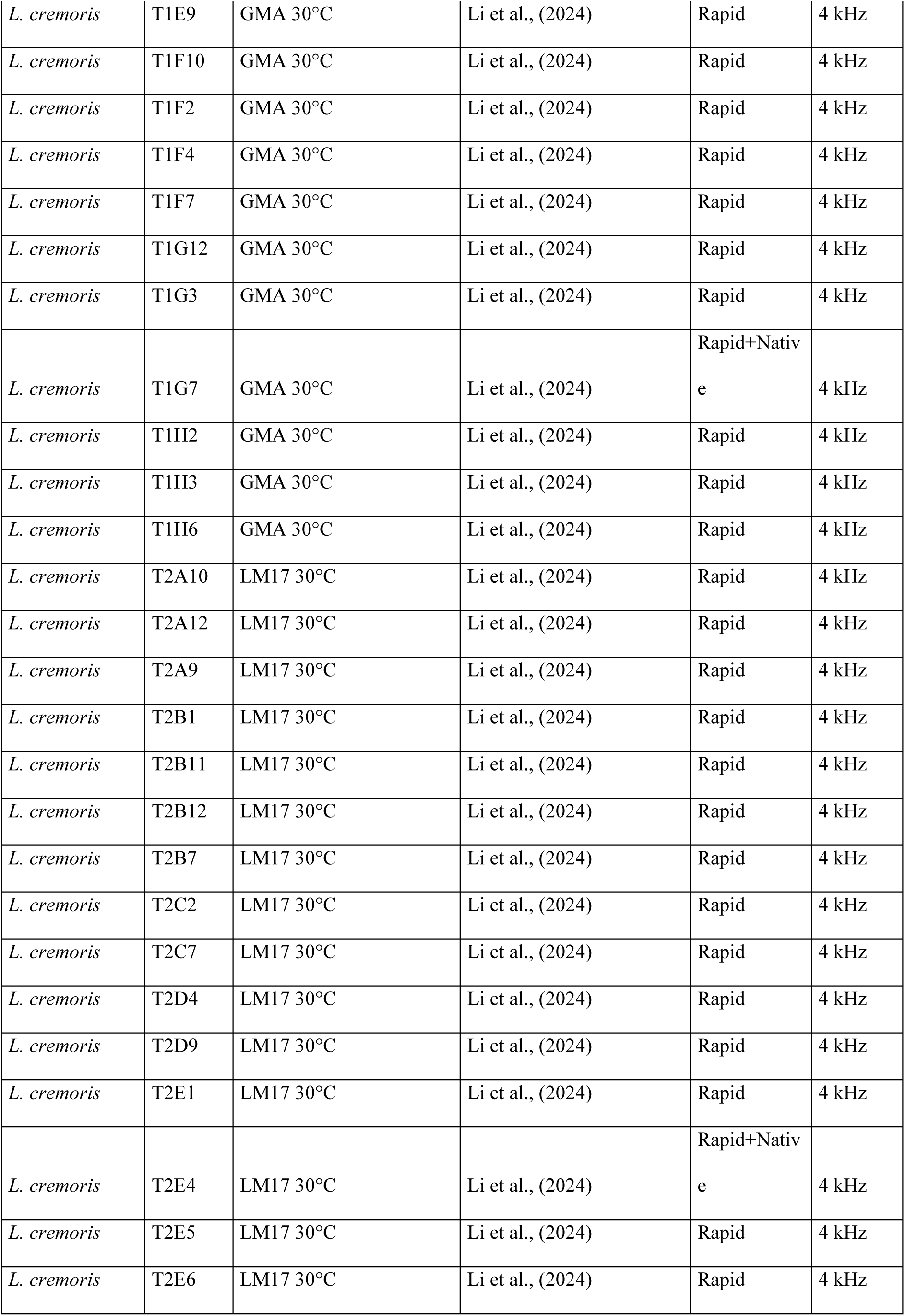
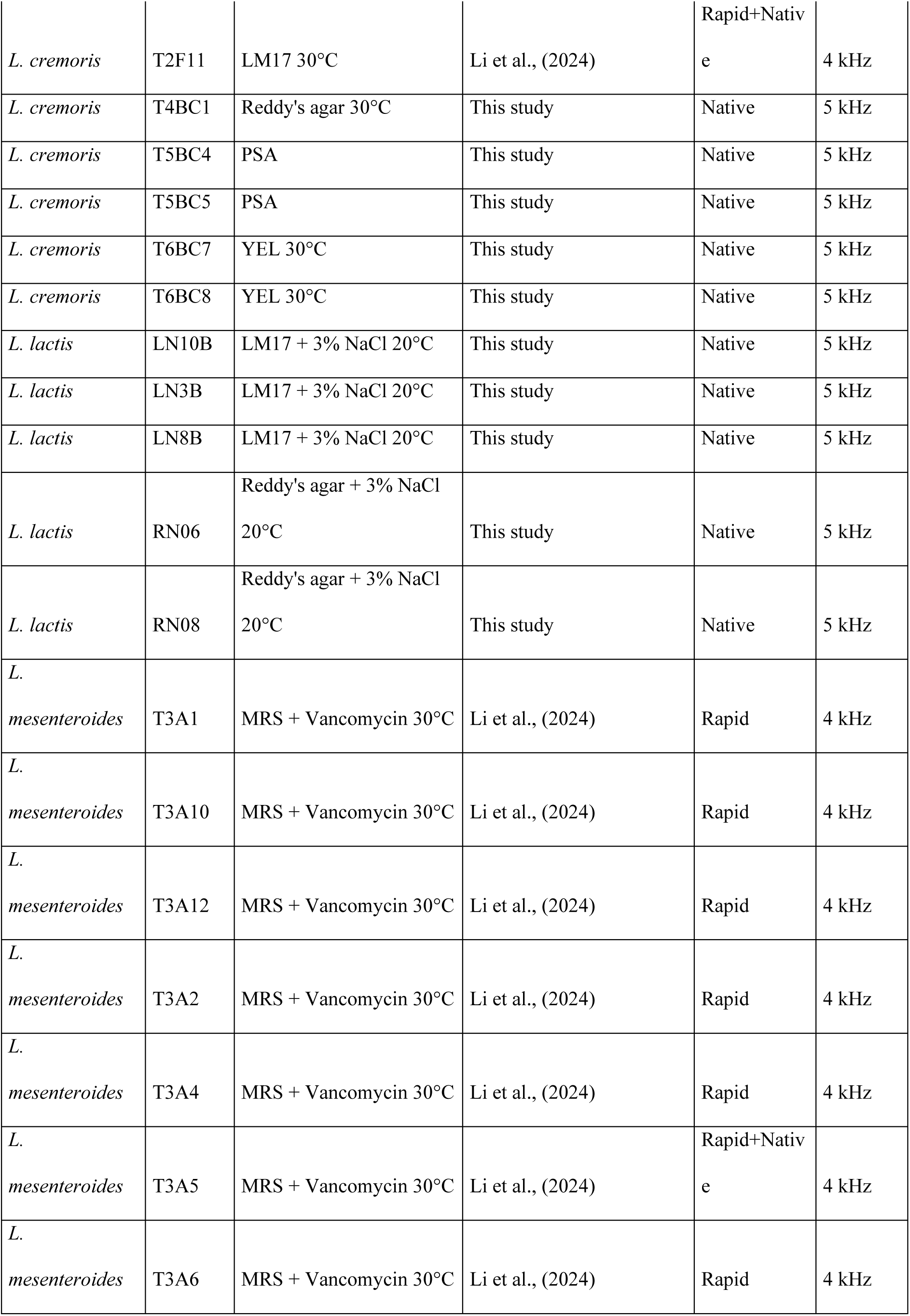
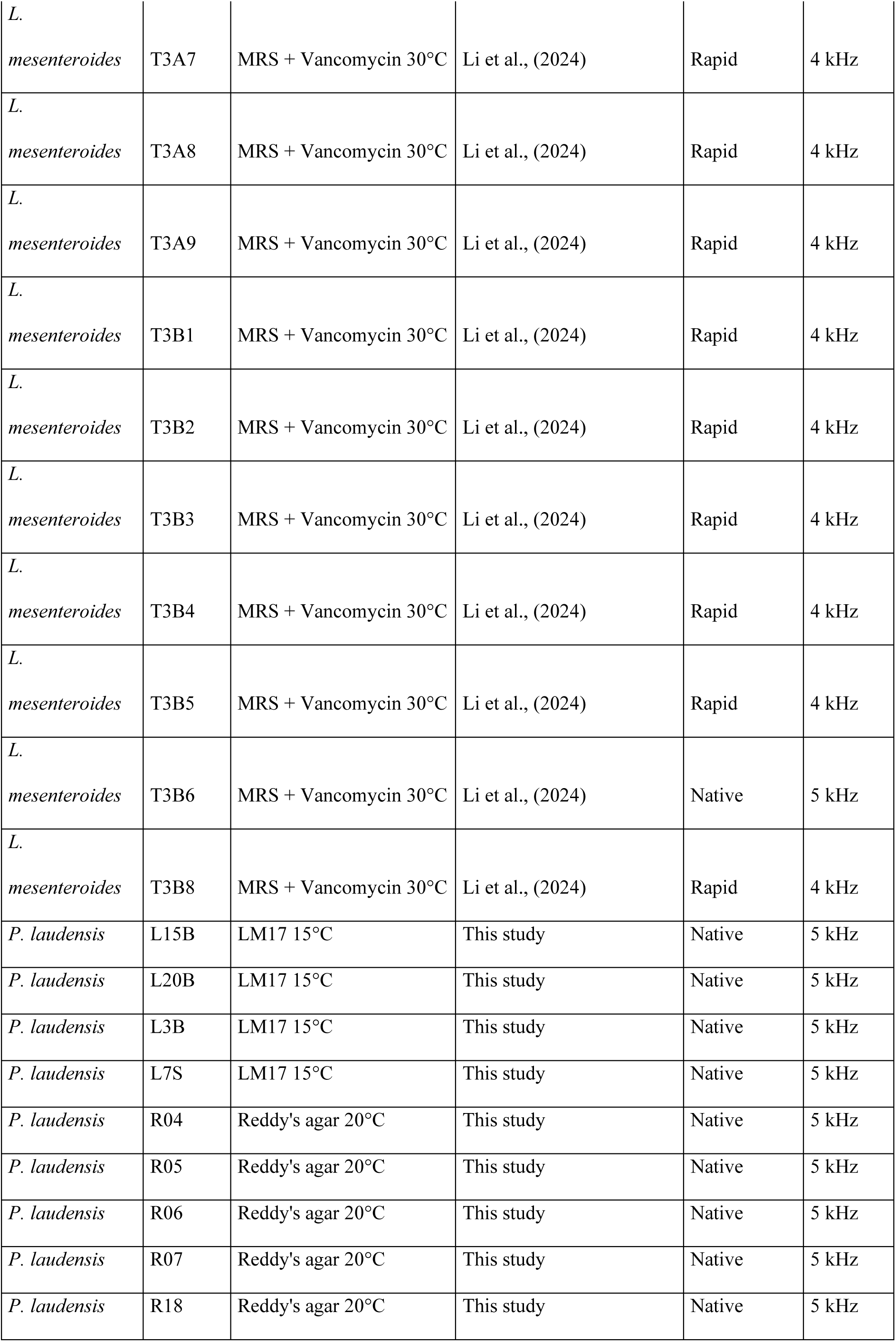
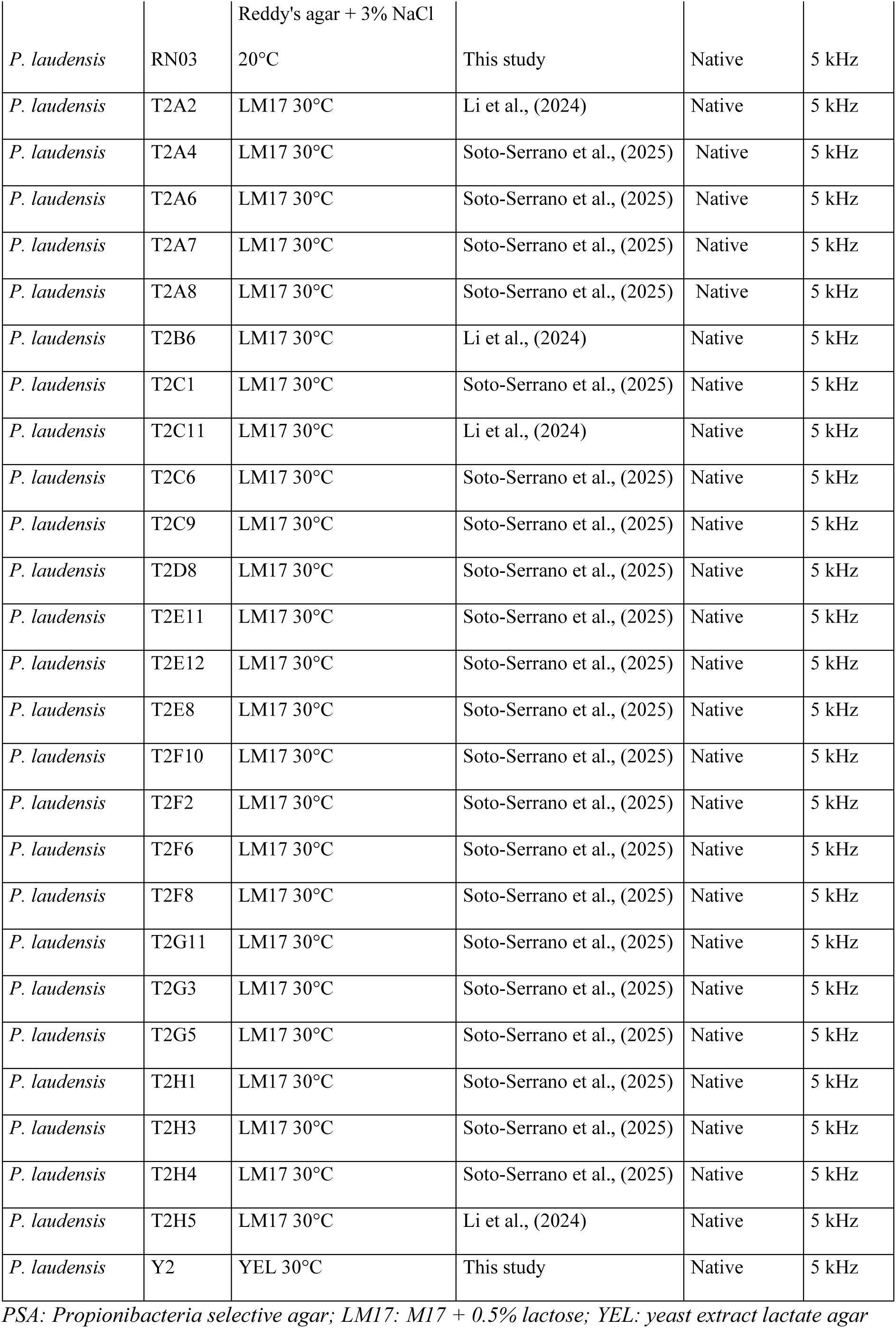
Strains used in this study.

**Sup. Fig. S1.**
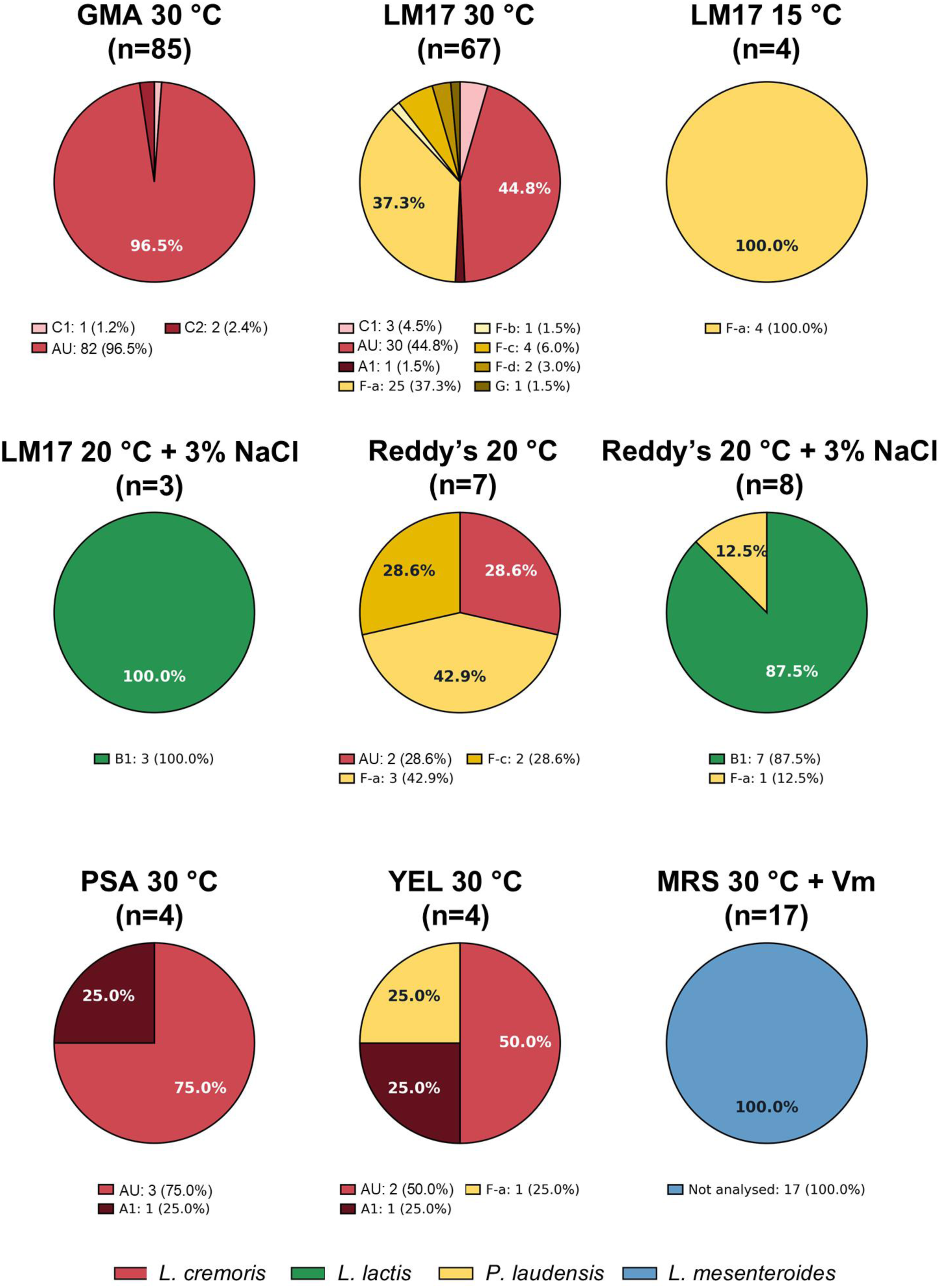
Species and *cwps* genotype representation across cultivation conditions. Pie charts show the composition of the isolate collection recovered under each medium, temperature and supplementation condition, before dereplication (199 isolates in total). Sector sizes indicate the proportion of isolates assigned to each cell wall polysaccharide (CWPS) genotype within a condition; legends report isolate counts and percentages, and n indicates the total number of isolates sampled per condition. Colours distinguish species: *L. cremoris* (red), *L. lactis* (green), *P. laudensis* (yellow) and *L. mesenteroides* (blue), with shades distinguishing CWPS genotypes within species. CWPS was not analysed in *L. mesenteroides*. Sampling intensity differed among conditions, with extensive initial sampling on GMA and LM17 at 30 °C and smaller subsequent collections targeting additional diversity. Vm, vancomycin.

**Sup. Fig. S2.**
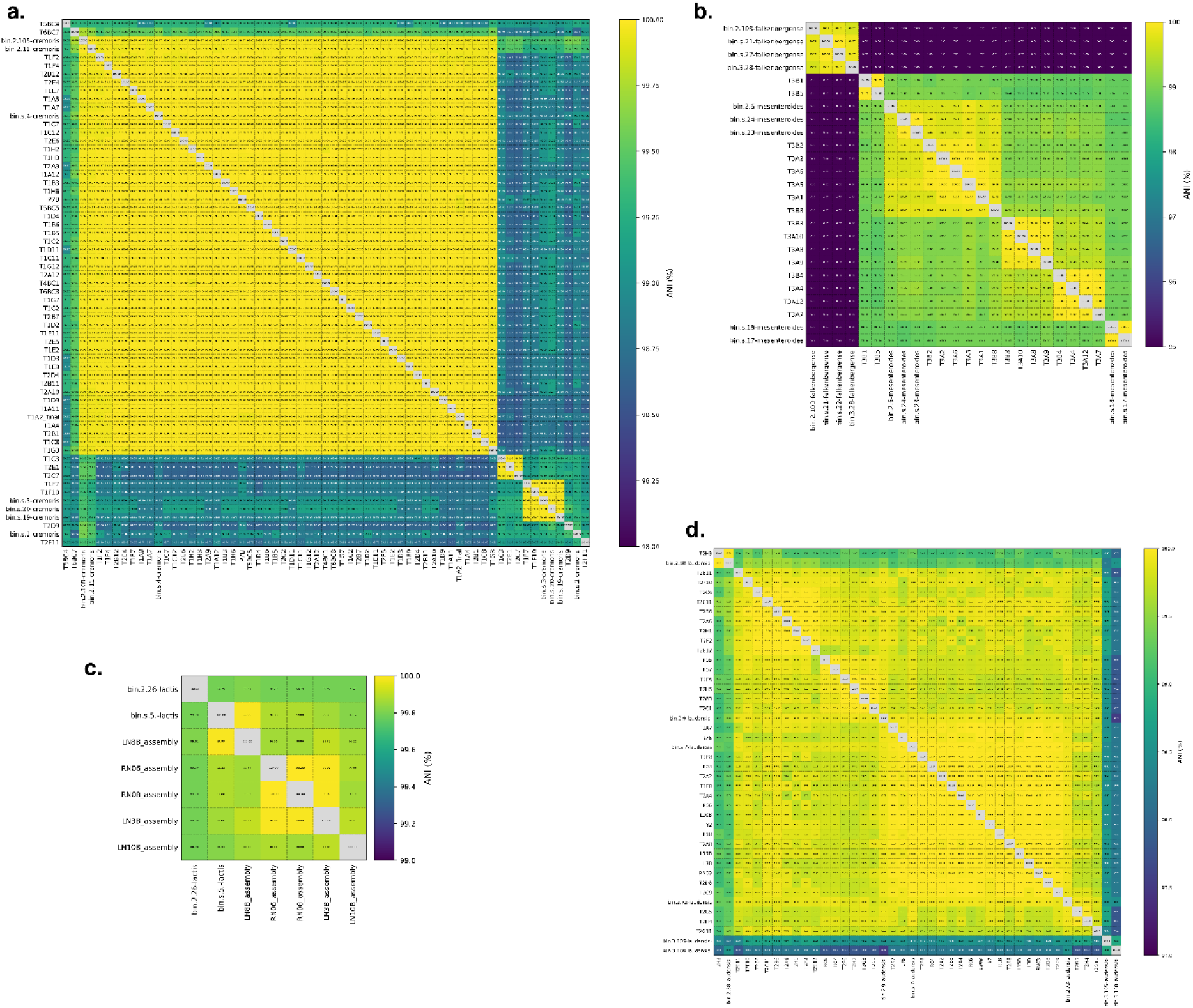
Average Nucleotide Identity between isolates and bins. (a) *L. cremoris*; (b) *Leuconostoc spp*.; (c) *L. lactis*; (d) *P. laudensis*

**Sup. Fig. S3.**
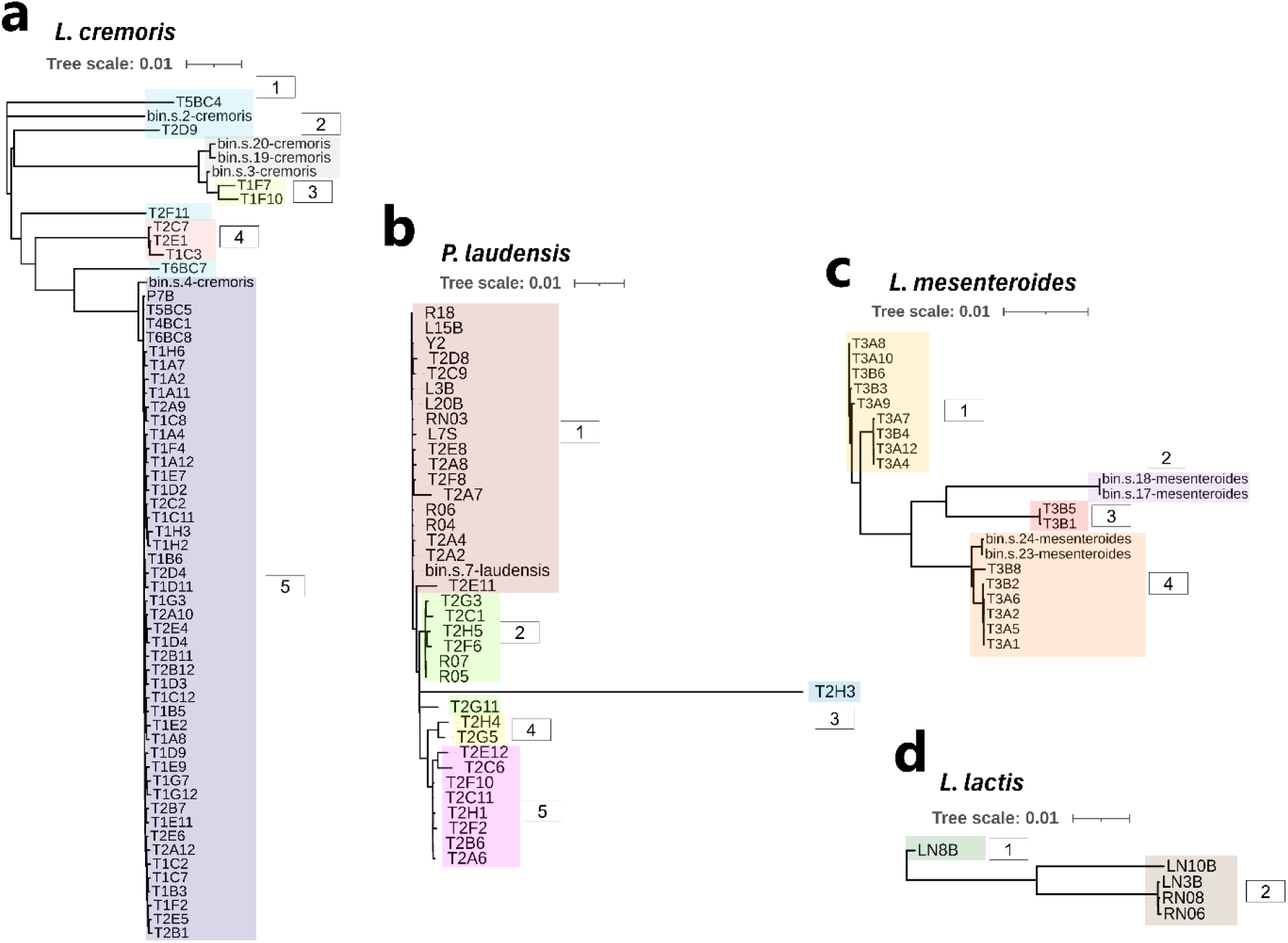
Core-genome phylogenies of isolate genomes and high-quality MAGs. Only MAGs with 0% strain heterogeneity. (labelled as “bin.*”) are included. (a) *L. cremoris*, (b) *P. laudensis*, (c) *L. mesenteroides*, and (d) *L. lactis*. Shaded clades denote within-sample lineages (numbers), illustrating partial overlap between isolate-derived and MAGs. Scale bars indicate substitutions per site.

**Sup. Fig. S4.**
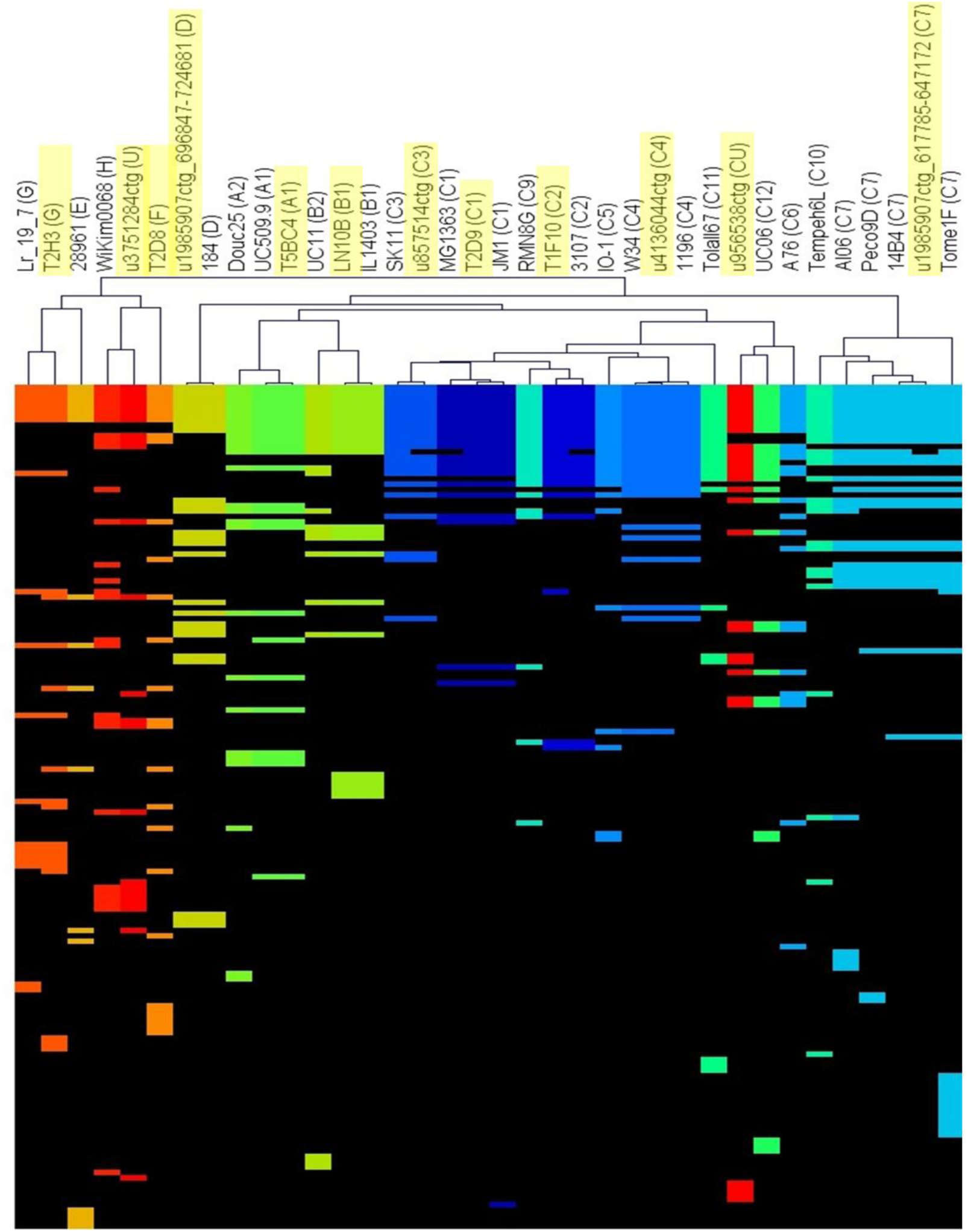
Two-way hierarchical clustering (HCL) of CWPS biosynthetic loci detected in isolates and metagenome contigs. Columns indicate representatives of *cwps*-containing genomes/contigs and rows represent genes; coloured tiles indicate presence of a given gene and black indicates absence, different colours represent the different genotypes, with highlighted genotypes corresponding to samples from the present study, while non-highlighted ones were used as genotype representatives.

**Sup. Fig. S5.**
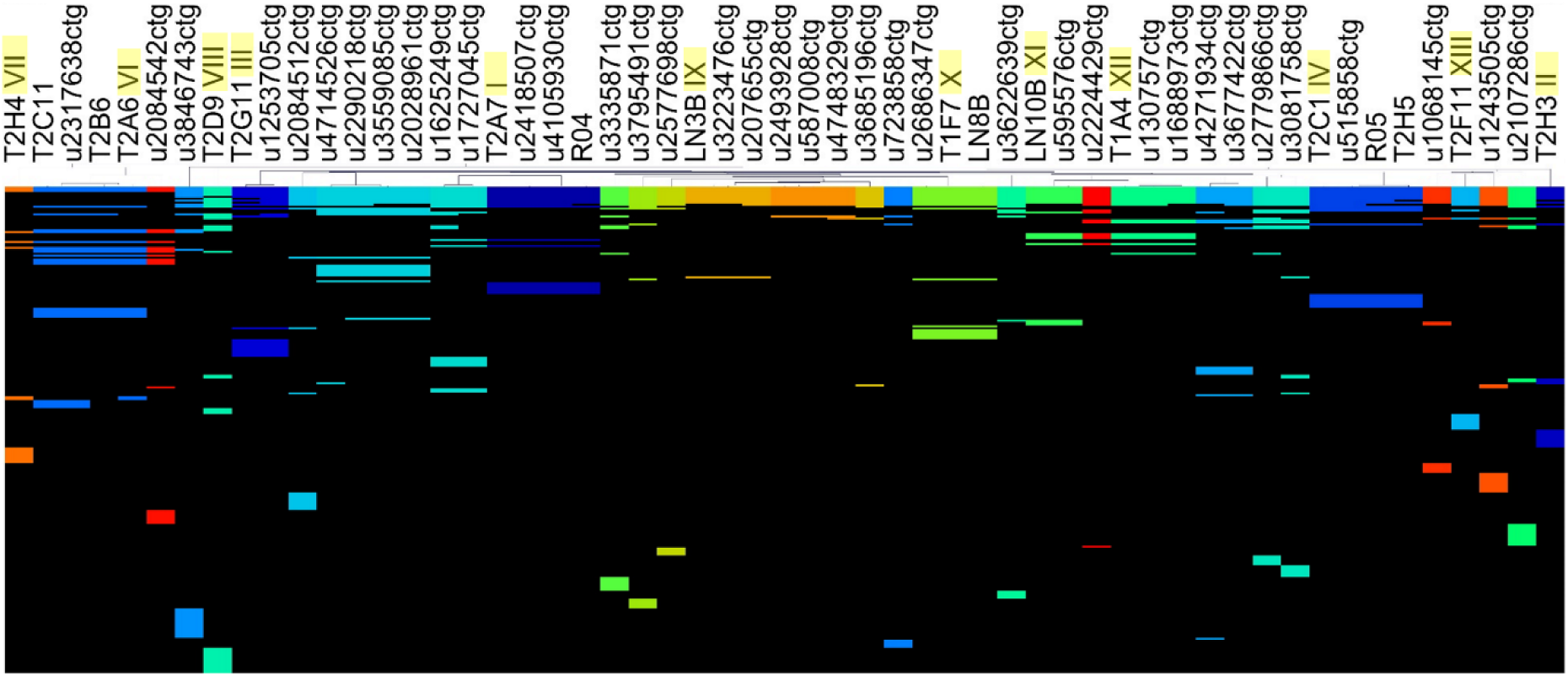
Two-way hierarchical clustering (HCL) of EPS biosynthetic loci detected in isolates and metagenome contigs. Columns indicate representatives of *eps*-containing genomes/contigs and rows represent genes; colored tiles indicate presence of a given gene and black indicates absence, different colors represent the different genotypes, with highlighted genotypes corresponding to isolate representatives (named I to XIV, where there is no genotype V for convention with a former study).

**Sup. Fig. S6.**
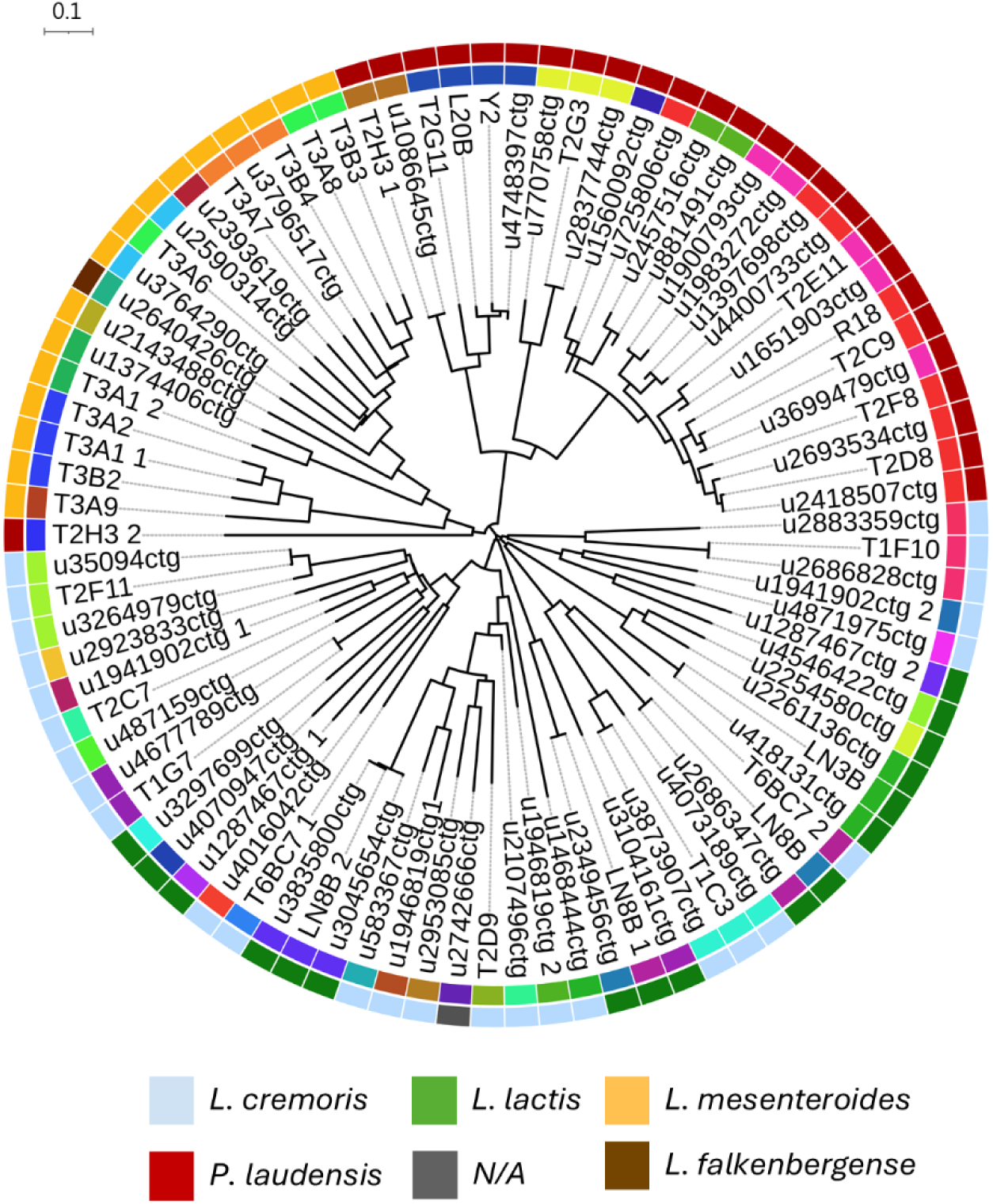
Proteome-based unrooted phylogeny of predicted intact prophages recovered from isolate genomes and metagenome contigs. The outer ring denotes the host species for isolate prophages or predicted host assignment for metagenome-derived prophages (legend), whereas the inner ring indicates prophage genus (49 genera total, each represented by a different color).

**Sup. Fig. S7.**
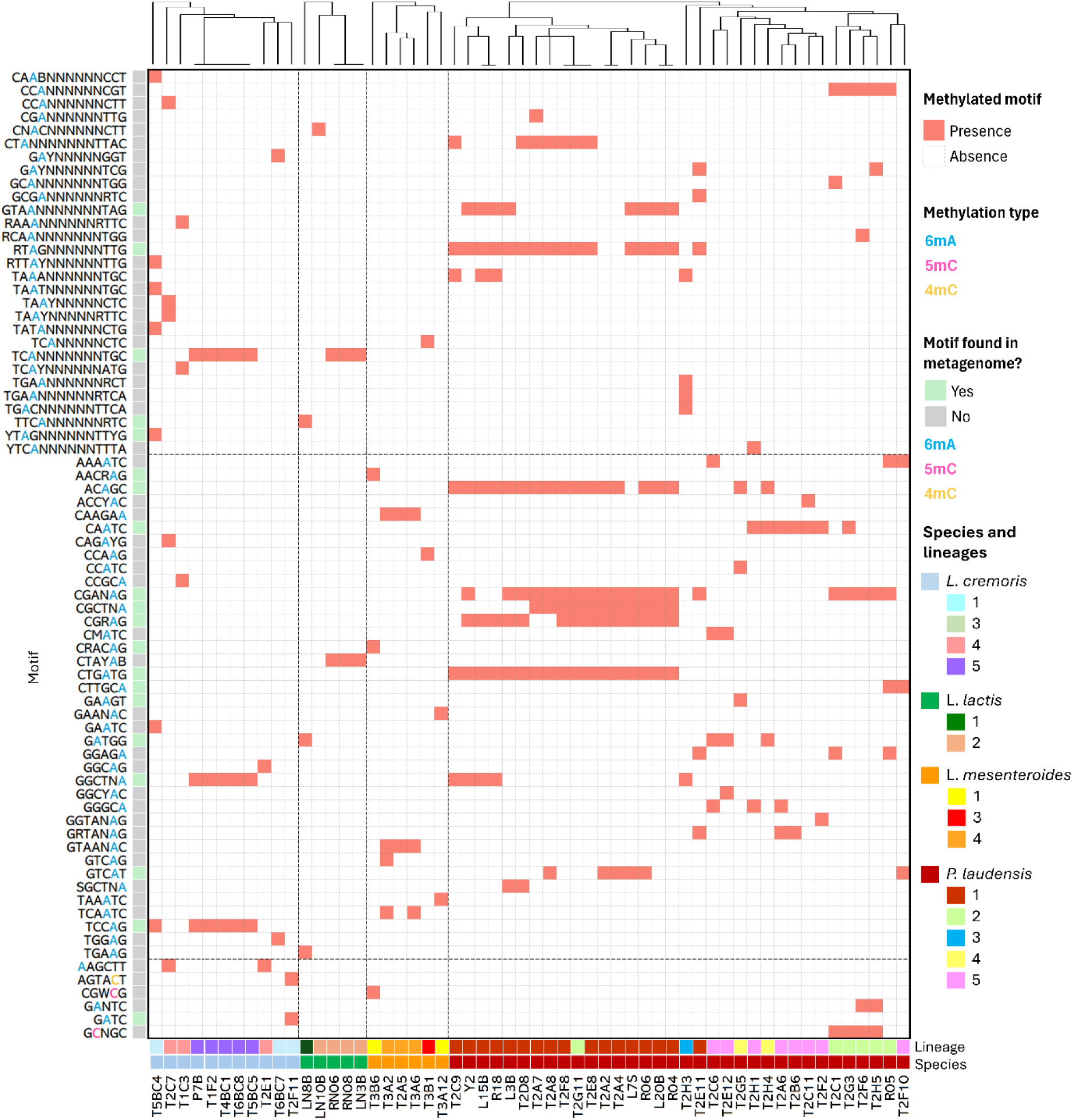
Isolate-derived strain-level methylomes. Heatmap showing the presence/absence of methylated DNA motifs across isolate genomes for which methylation calling was available. Rows represent distinct methylated motifs, with coloured nucleotides denoting methylation type (6mA, 5mC or 4mC). Columns represent individual isolates, hierarchically clustered by motif repertoire similarity within species (top). Motifs also detected in the metagenome dataset are indicated (legend). Second annotation row represents lineages. Vertical dashed separators indicate species groupings, while horizontal dashed separators indicate motif type: bipartite (top), non-palindromic (middle), palindromic (bottom).

**Sup. Fig. S8.**
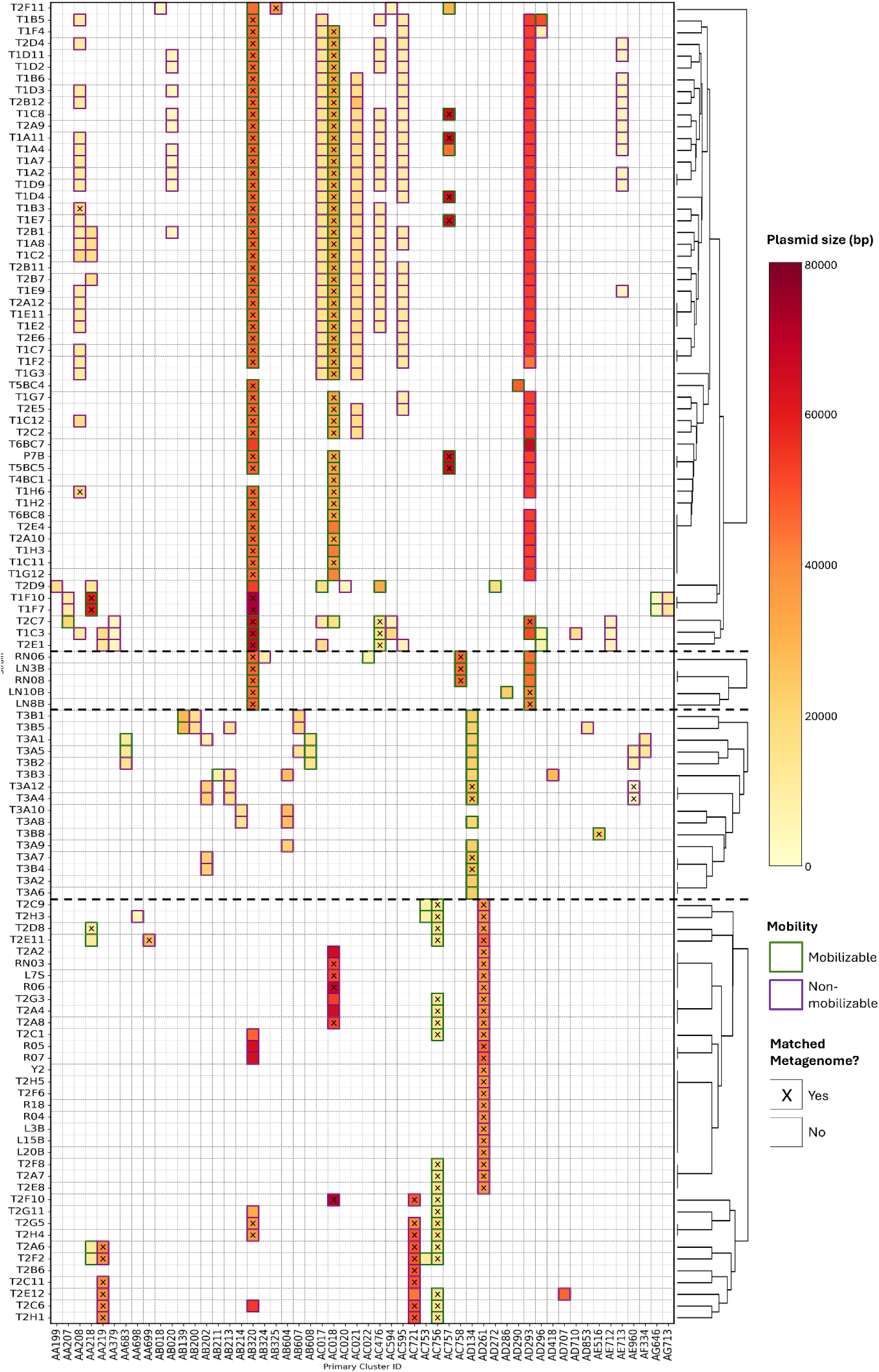
Plasmidome composition of the bacterial isolates, mobility potential, and overlap with metagenome-assembled circular plasmids. The heatmap shows the distribution of circular plasmids across the non-redundant isolate collection (rows) grouped by host species (dashed separators, from top to bottom: *L. cremoris, L. lactis, L. mesenteroides, P. laudensis*). Columns correspond to MOB-suite primary plasmid cluster IDs. Each coloured cell indicates the presence of a plasmid in each isolate, with colour intensity representing plasmid length (bp). Plasmid mobility potential as predicted by MOB-suite is indicated by the cell outline (mobilizable versus non-mobilizable). Plasmids also recovered as circular contigs in the metagenome assembly are marked with an “X”. Hierarchical clustering (right) indicates similarity in plasmid composition across isolates from the same species.

**Sup. Fig. S9.**
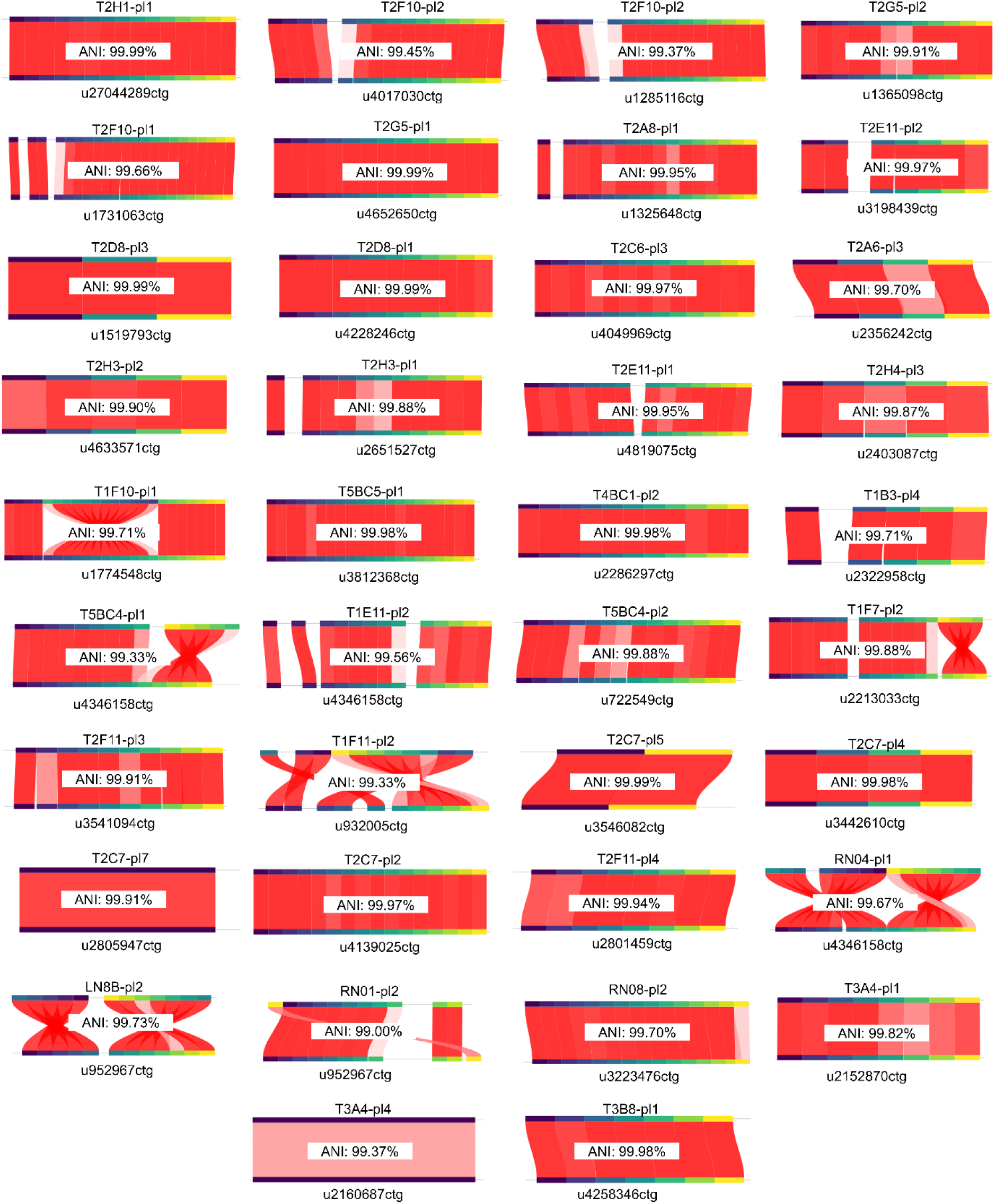
Sequence comparisons between plasmids recovered from isolates and metagenomics. Pairwise comparisons were generated using FastANI, and show isolate-derived plasmids above and circular metagenomic plasmid contigs below, with their corresponding identifiers. Red connections represent reciprocally mapped sequence fragments and illustrate their relative positions and orientations. ANI estimates are indicated within each panel. Only comparisons with ANI ≥99.0% and aligned fraction ≥80.0% were considered.

**Sup. Fig. S10.**
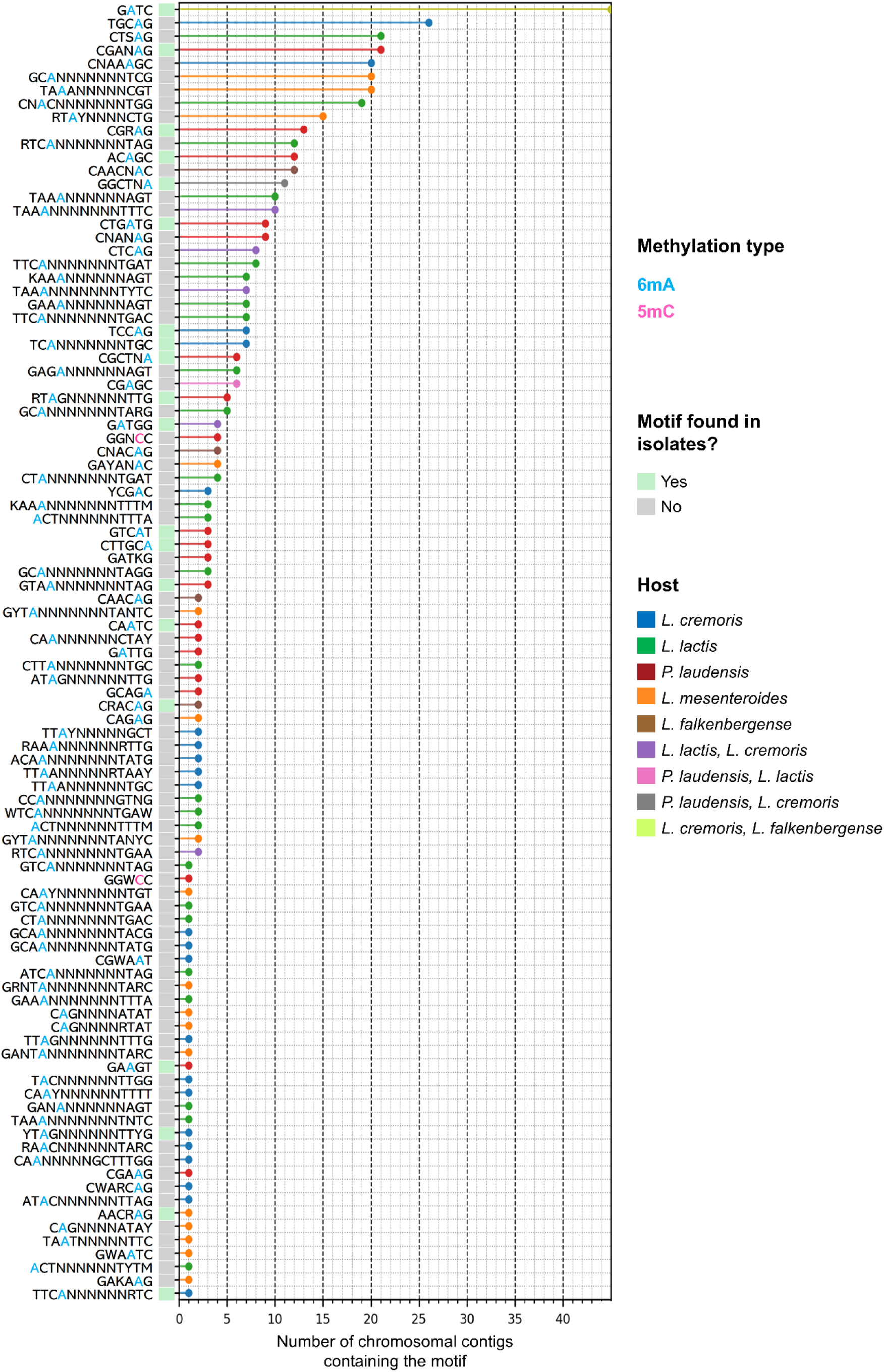
Methylated DNA motifs identified from metagenome-assembled chromosomal-only contigs. Rows indicate individual motifs after manual curation of duplicated and ambiguous motif calls, with coloured nucleotides representing methylation type (6mA, 5mC or 4mC). Coloured cells indicate motifs that have been also found in the isolates set. The x-axis denotes the number of chromosomal contigs carrying each motif, while the taxonomically assigned host(s) to the chromosomal contigs containing the motifs is indicated in the colour bars.

